# Small-Molecule Modulators of Lipid Raft Stability and Protein-Raft Partitioning

**DOI:** 10.1101/2024.10.28.620521

**Authors:** Katherine M. Stefanski, Hui Huang, Dustin D. Luu, James M. Hutchison, Nilabh Saksena, Alexander J. Fisch, Thomas P. Hasaka, Joshua A. Bauer, Anne K. Kenworthy, Wade D. Van Horn, Charles R. Sanders

## Abstract

Development of an understanding of membrane nanodomains colloquially known as “lipid rafts” has been hindered by a lack of pharmacological tools to manipulate rafts and protein affinity for rafts. We screened 24,000 small molecules for modulators of the affinity of peripheral myelin protein 22 (PMP22) for rafts in giant plasma membrane vesicles (GPMVs). Hits were counter-screened against another raft protein, MAL, and tested for impact on raft, leading to two classes of compounds. Class I molecules altered the raft affinity of PMP22 and MAL and also reduced raft formation in a protein-dependent manner. Class II molecules modulated raft formation in a protein-independent manner. This suggests independent forces work collectively to stabilize lipid rafts. Both classes of compounds altered membrane fluidity in cells and modulated TRPM8 channel function. These compounds provide new tools for probing lipid raft function in cells and for furthering our understanding of raft biophysics.

**Teaser:** Compounds have been discovered that modulate the affinity of membrane proteins for lipid rafts as well as raft formation.

## Introduction

Lipid rafts are dynamic phase-separated nanodomains of the plasma membrane that are thicker and more rigid than the bulk membrane and are typically enriched in cholesterol and in phospholipids with saturated acyl chains, often sphingolipids (1–4). It has long been theorized that cells utilize lipid rafts as sorting and signaling platforms (5–10). Specific proteins are thought to partition into rafts under cellular conditions where they facilitate a variety of biological functions. Lipid rafts have been implicated in immune signaling, neural development, host-pathogen interactions, caveolae, cytoskeletal-membrane contacts, and other cellular phenomena (11–15). Under physiological conditions lipid rafts in cells are believed to be nanoscale and highly dynamic, severely limiting traditional microscopic studies (16–18).

Giant plasma membrane vesicles (GPMVs) derived from the plasma membranes of live cells are a well-established tool for studying lipid rafts and for quantitating the raft affinity of membrane proteins (19). GPMVs are easy to prepare and will spontaneously phase separate into micron-scale liquid ordered (L_o_, raft) and liquid disordered (L_d_, non-raft) domains. Assorted fluorescent dyes or protein markers are available to label ordered and disordered phases in GPMVs, enabling imaging-based analysis of phases and resident proteins. GPMVs also retain the lipid and protein composition of the cell plasma membrane from which they are derived (20, 21).

The affinity of a number of membrane proteins for lipid rafts relative to the surrounding disordered phase has been assessed in GPMVs. GPMVs have also been used to elucidate the characteristics that drive raft affinity (22–24). A majority of the membrane proteins thought to preferentially partition into lipid rafts are single-span proteins. Studies of single-span raft-avid proteins enabled the identification of structural criteria for raft affinity. Palmitoylation, transmembrane domain length, and surface area are the main drivers of raft partitioning for single-pass membrane proteins (22). No such rules have been determined for multispan proteins, in large part because only a small number of raft-favoring multispan proteins have been unambiguously identified, and the rules for single-pass proteins do not seem to apply (23, 25).

Peripheral myelin protein 22 (PMP22) is a tetraspan membrane protein that preferentially partitions into ordered domains in GPMVs. PMP22 is highly expressed in myelinating Schwann cells and is a major component of myelin in the peripheral nervous system (PNS). Duplication of the PMP22 gene causes the most common type 1A form of Charcot-Marie-Tooth (CMT) disease (26), a peripheral neuropathy that is a top 10 most common genetic disorder. More severe forms of CMT are caused by inherited point mutations in PMP22 (27). While the biophysical properties that drive PMP22 into lipid rafts are unknown, an analysis of disease-associated PMP22 mutations in GPMVs revealed that most of the mutations resulted in reduced protein preference for the raft phase versus the disordered phase (25). The fact that the tested disease mutants are known to destabilize protein folding suggests raft affinity is dependent on the fold of PMP22 (28). Given that PMP22 is thought to be involved in supporting cholesterol homeostasis and trafficking in Schwann cells, it is very possible that its affinity for lipid rafts is closely related to one of its key physiological functions in forming healthy myelin (29). These considerations motivated us to seek compounds that alter PMP22 raft affinity, thereby providing tools for use in investigating the drivers and physiological consequences of PMP22 raft partitioning.

We conducted high throughput screening (HTS) to discover molecules that alter PMP22 phase partitioning between ordered and disordered domains. Our primary assay reports not only on PMP22 phase partitioning, but also on whether compounds alter raft formation. Here we report the discovery of small molecules falling into two classes. Class I compounds decreased the affinity for lipid rafts of PMP22 and myelin and lymphocyte protein (MAL) [another raft-preferring tetraspan membrane protein (23)]. Class I compounds also reduced raft formation in a protein-dependent manner. Class II compounds altered raft formation independent of proteins. Both classes of compounds were seen to alter membrane fluidity in GPMVs and live cells. We further explored their activities by examining their effects on the activity of the human cold-sensing TRPM8 channel and on signaling by the epidermal growth factor receptor (EGFR). These compounds provide new tools for investigating raft-dependent phenomena in cells. Additionally, the differing modes of action of these compounds expand our understanding of raft biophysics.

## Results

### A high-throughput screen identifies compounds that alter PMP22 raft affinity

To identify compounds that alter the ordered domain partitioning of PMP22 we conducted a high-throughput screen adapted from our recent method (Fig. 1) (25). Briefly, the goal was to use GPMVs from cells expressing PMP22 to screen for small molecules that alter the proportion of PMP22 in the ordered phase. GPMVs were made from transfected HeLa cells expressing PMP22. We employed the N41Q (glycosylation deficient) variant of PMP22 since it traffics to the plasma membrane with much higher efficiency than wild type (WT) but exhibits the same ordered domain preference (25, 31). The disordered phase was labeled with the lipophilic stain DiI while PMP22 was immunolabeled with AlexaFluor 647. GPMVs were deposited into 96 or 384-well plates containing compounds at a final concentration of 10 µM. A 23,360 compound sub-set of the >100,000 compound Vanderbilt Discovery Collection and a library of FDA approved drugs (1,184) were screened. The Discovery collection compounds represents a cross-section in terms of the chemical diversity of the master library. Plates were imaged using a high-content spinning disc confocal imaging system. VesA software was used to analyze images (30). The power of this approach is in the number of vesicles that can be imaged and analyzed in an unbiased manner. A typical well results in images containing thousands of GPMVs which are then analyzed automatically, independent of investigator bias or foreknowledge. Hits from the screen were picked using strictly standardized mean difference (SSMD) values on a plate-by-plate basis (typical cutoff was an SSMD value ≥ 90% of the positive control) based on compound impact on PMP22 *P_ordered_* (32). *P_ordered_* is the fraction of protein in the ordered phase and is a measure of the affinity of a particular protein for rafts. An initial hit rate of 1.06% was observed, resulting in 267 preliminary hits. Hits were then confirmed in triplicate experiments using compounds from the library. Compounds of interest were then reordered from the manufacturer and their effects confirmed. Validated compounds were then tested in GPMVs from cells expressing the MAL protein, another lipid raft-preferring tetraspan protein (23), to test whether their activity is protein-specific.

**Figure 1.**
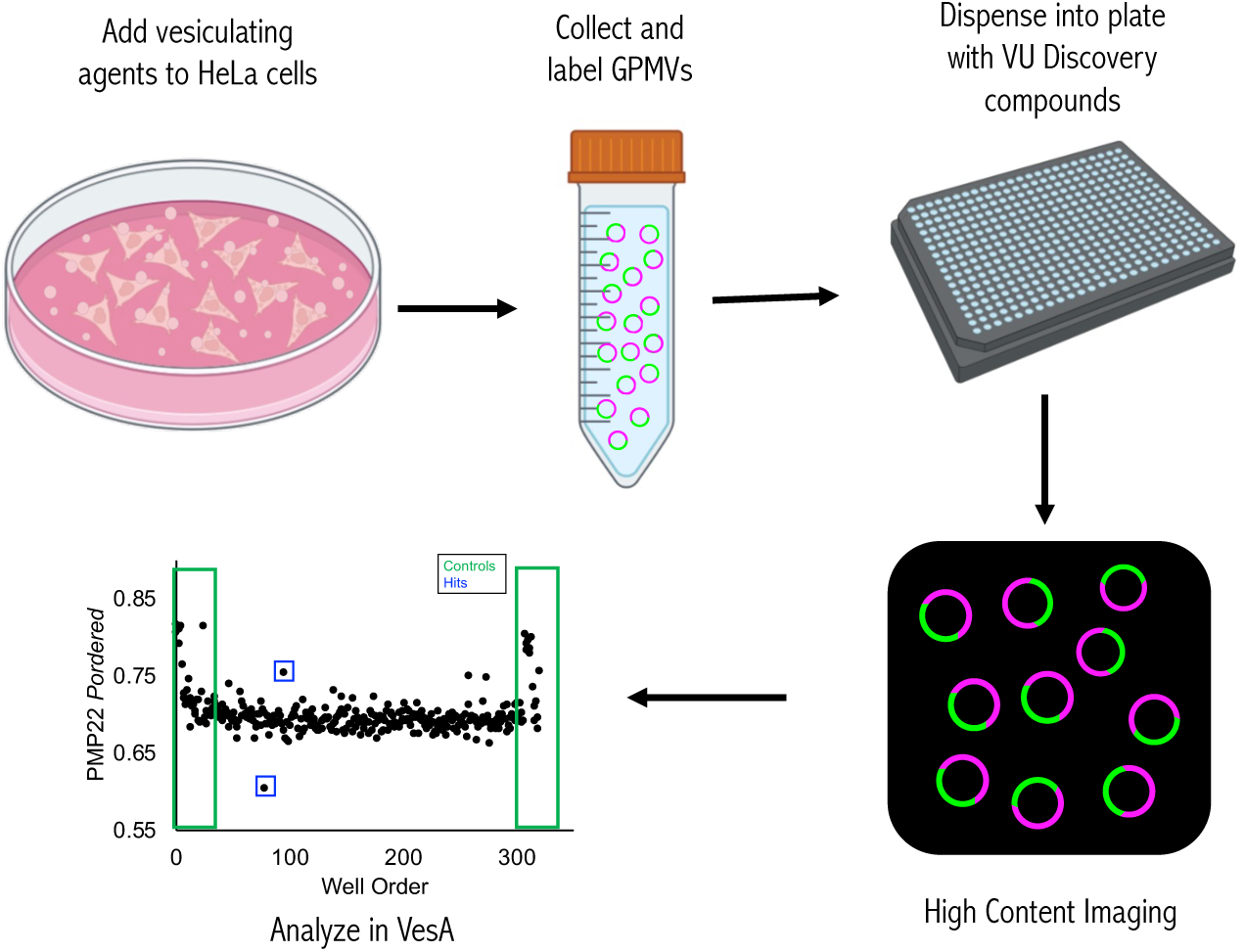
High-throughput screening approach to identify modulators of PMP22 raft affinity. Pipeline used to screen 24,000+ compounds that identified hits described here.

Effects of all hits on raft formation were also measured. Here we used the fraction of phase-separated GPMVs (vs non phase-separated) in our images as a measure of raft formation. For all confirmed compound hits (above), we also completed measurements in GPMVs from untransfected cells. Hits fell into several distinct categories based on these data criteria. In this work, we focus on two interesting functional classes of compounds.

### Protein-dependent modulators of raft affinity and raft stability (Class I molecules)

Among the validated hits, two compounds were found to have unique effects on both protein raft affinity and raft formation. VU0615562 and VU0619195 significantly decreased *P_ordered_* for both PMP22 and MAL by ∼18-20% (Figs. 2A and 2B). While PMP22 and MAL are tetraspan myelin proteins with a known affinity for lipid rafts they have little-or-no sequence homology. These compounds thus reduce *P_ordered_* for two non-homologous tetraspan membrane proteins.

**Figure 2.**
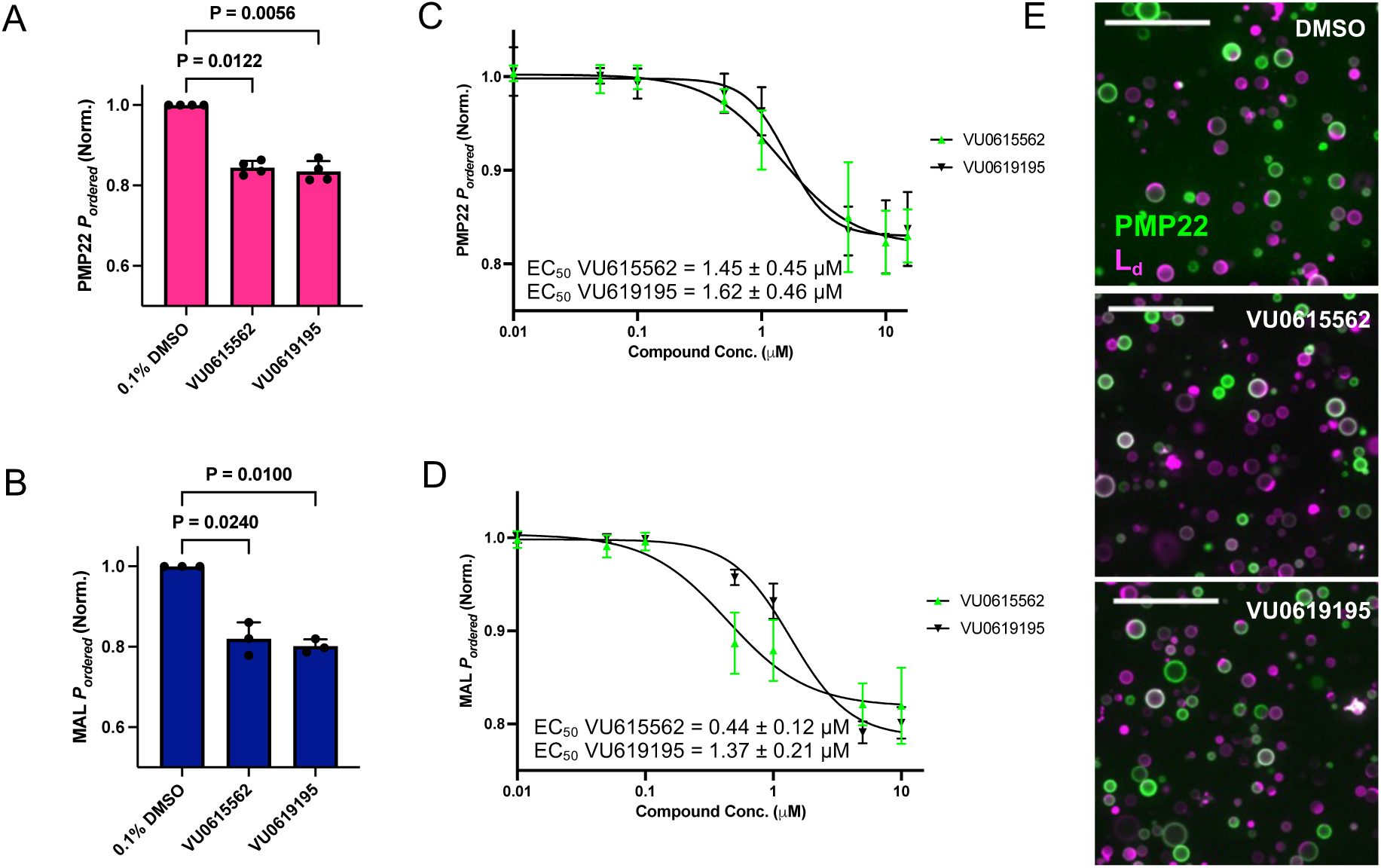
Class I modulators affect the affinity of PMP22 and MAL for ordered domains. **A)** Hit compounds VU0615562 and VU0619195 decrease ordered partitioning of PMP22 (*n* = 4) and **B)** MAL (*n* = 3) at 10 μM, Bars are means ± SD. P-values are from Mann-Whitney tests. **C)** Dose-response experiments determined EC_50_ for the impact of VU0615562 and VU0619195 on PMP22 ordered partitioning and **D)** MAL ordered partitioning. Points are mean ± SD, *n* = 3. Curves and EC_50_ values are from non-linear, sigmoidal fits, ± SE. **E)** Representative images of GPMVs treated with 0.5 μM compound. Scale bars are 50 μm.

Dose-response experiments indicated that EC_50_ values for the impact of the two compounds on each protein are in the ∼1 µM range (Figs. 2C and 2D). To verify the effects of the compounds are not specific to HeLa cells we confirmed similar activities in GPMVs derived from rat basophilic leukemia (RBL) cells which also produce GPMVs that phase separate at temperatures that are feasible for our microscopes (fig. S1).

We next examined the effects of VU0615562 and VU0619195 on raft stability in GPMVs derived from untransfected cells and from cells expressing either PMP22 or MAL. We used the fraction of phase-separated vesicles from images (c.f., Fig. 2E) as a proxy for raft formation. The compounds decreased the fraction of phase-separated GPMVs in all cases (Figs. 3A and 3B, gray bars). However, when PMP22 or MAL were overexpressed, this reduction in phase-separated vesicles was significantly more pronounced compared with GPMVs from untransfected cells (Figs. 3A and 3B) and was observed across a range of concentrations (Fig. 3C, curve does not approach 0 as in Figs. 3D and 3E). For both compounds the EC_50_ values were in the vicinity of 1 µM. This led us to speculate that these compounds reduce raft stability via a mechanism that is not specific to a single protein but is dependent on the presence of proteins. We hypothesize that the activity of the compounds observed in GPMVs from untransfected cells is because the compounds exploit endogenous membrane proteins in HeLa cells to exert their effects.

**Figure 3.**
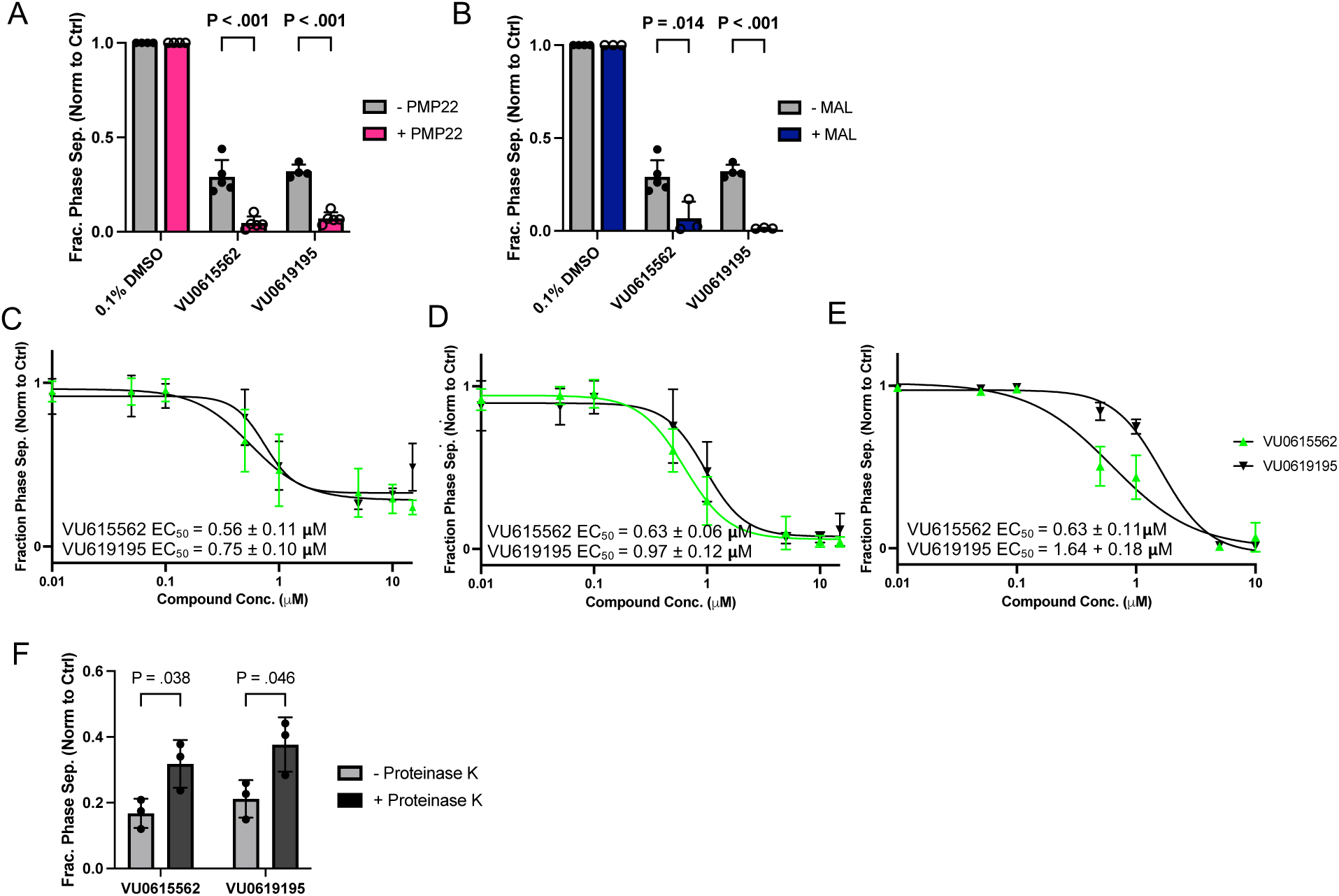
Class I compounds decrease raft formation in a protein-dependent manner. **A)** Effects of 10 μM VU0615562 and VU0619195 on the fraction of phase separated GPMVs with PMP22 expression (magenta bars) or from untransfected cells (gray bars) (*n* = 4-5). **B)** Effects of VU0615562 and VU0619195 on the fraction of phase separated GPMVs with MAL expression (navy bars) or from untransfected cells (gray bars) expression 10 μM (*n* =3-5). **The middle three panels present dose response experiments** used to determine EC_50_ values of compounds on phase separation in GPMVs from **C)** cells not expressing PMP22 or MAL, **D)** cells expressing PMP22, and **E)** cells expressing MAL. *n* = 3, points are means ± SD. Curves and EC_50_ values ± SE are from non-linear, sigmoidal fits. **F)** Effects of 10 μM compounds on the fraction of phase separated GPMVs with (dark gray bars) and without proteinase K treatment from untransfected cells (*n* = 3). Bars are means ± SD. P-values are from unpaired student’s t-tests.

We know that PMP22 expression stabilizes ordered domains (25) so the augmented effect size of raft destabilization observed in GPMVs + PMP22 seems to be the consequence of an intrinsic property of VU0615562 and VU0619195 that is somehow enhanced when PMP22 is present. To determine if MAL has a similar effect on raft formation in the absence of Class I compounds we used data from within the same images and compared the fraction of phase-separated vesicles from MAL positive (GFP+) and MAL negative (GFP-) GPMVs (a feature of the VesA software allows us to make this distinction). We used DMSO treated control wells from dose-response experiments for this comparison. In contrast to PMP22 (25), in compound-free samples we found that MAL expression has very little impact on phase separation (fig. S2). These data indicate that the enhancement of Class I compound raft destabilization by PMP22 or MAL reflects an intrinsic activity of these compounds that is enhanced by the presence of the proteins via an unknown mechanism.

To further investigate this apparent protein dependence, we tested the impact of these compounds on GPMVs treated with proteinase K. Proteinase K is a broad-spectrum serine protease capable of cleaving proteins preferentially after hydrophobic residues. Note that it has been previously shown that GPMVs can be porous, so it is likely both intracellular and extracellular loops and segments are cleaved by the protease (33). We applied this treatment to GPMVs from untransfected cells because the protease activity would eliminate our ability to label or detect PMP22 or MAL (by cleaving the myc tag or GFP fusion), obviating our ability to differentiate GPMVs containing either protein. We found that proteinase K treatment significantly reduced the effects of the compounds (Fig. 3F). This supports the conjecture that the changes induced by the compounds are not specific to any one protein but are dependent on the presence of multiple proteins (overexpressed and/or endogenous). The incomplete effect of proteinase K in reducing the raft-lowering activities of these compounds may be due to its inability to digest all membrane proteins present completely. However, these data, combined with the results seen with PMP22 and MAL led us to conclude that multiple proteins have the trait of being able to enhance the ordered phase destabilizing effects of VU0615562 and VU0619195.

To be certain that the compounds were not remodeling the GPMVs we examined the size of GPMVs and the size of the ordered phases when treated with the compounds. Neither GPMV radii nor relative sizes of ordered phase domains were changed by VU0615562 or VU0619195 (figs. S3A and S3B), even though they decrease the raft affinities of PMP22 and MAL, as well as raft formation.

An examination of the chemical structures of VU0615562 and VU0619195 (Table 1) reveals a Tanimoto coefficient (a metric of chemical similarity ranging from 0-1) of 0.7778 (34). This chemical similarity is unsurprising in light of their similar effects on *P_ordered_* for PMP22 and MAL, as well as raft formation. Since these compounds were similar, structure-activity relationship (SAR) studies were conducted on chemically similar compounds available in the VU Discovery Collection (fig. S4). For VU0615562, seven additional compounds with similar structures (similarity integer value of 97) were tested. For VU0619195, six additional compounds with similar structures (similarity integer value of 85) were tested. None of the compounds tested had a greater impact on PMP22 ordered partitioning or raft stability than VU0615562 and VU0619195 (fig. S4). It is interesting, however, that for both compounds, analogs were found that maintained the capacity of the parent compound to reduce the raft affinity of PMP22 while largely losing their ability to reduce raft formation, suggesting that these two activities are incompletely coupled.

**Table 1.**
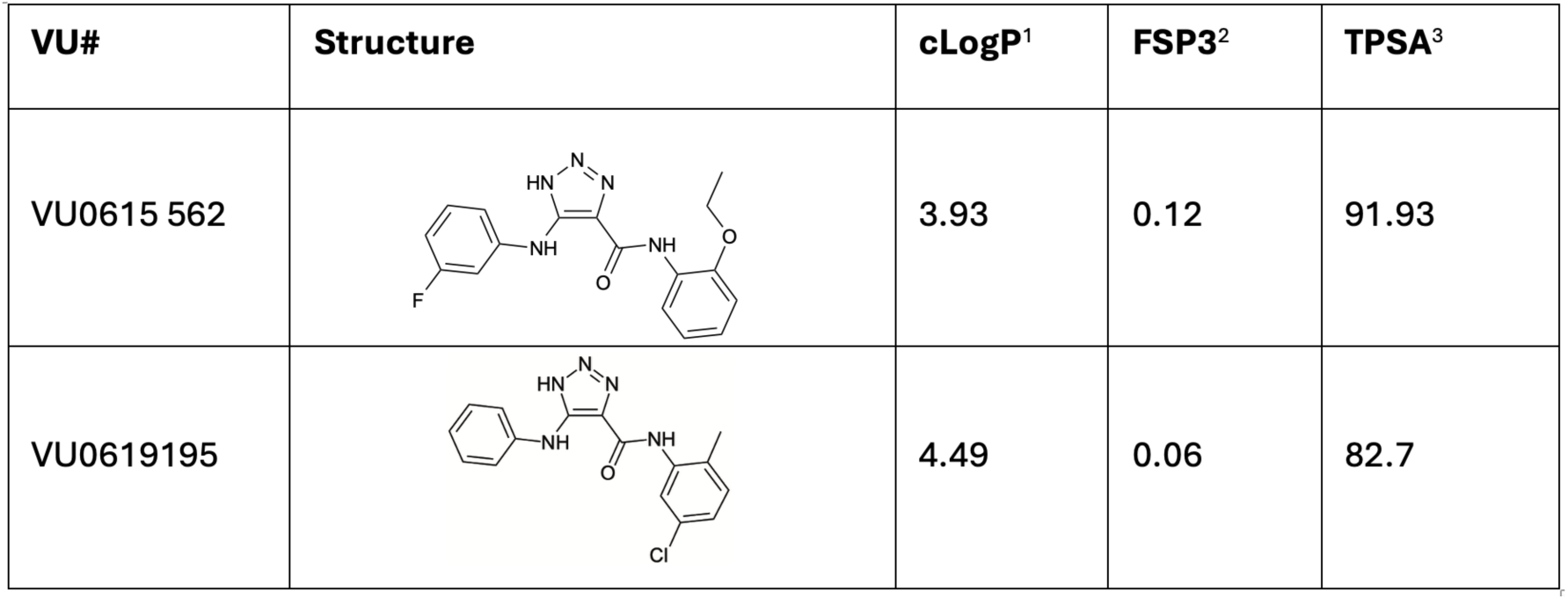
Protein-dependent compound structures and parameters.

| VU# | Structure | cLogP<sup>1</sup> | FSP3<sup>2</sup> | TPSA<sup>3</sup> |
| --- | --- | --- | --- | --- |
| VU0615 562 |  | 3.93 | 0.12 | 91.93 |
| VU0619195 |  | 4.49 | 0.06 | 82.7 |

### Protein-independent raft modulators (Class II)

It is reasonable to wonder if the protein-dependent raft modulating effects described above for VU0615562 and VU0619195 are unique to these compounds or if all raft modulating compounds would show a similar dependence in their activities on proteins. This turned out not to be the case based for three other compounds discovered in the initial screen for compounds that altered *P_ordered_* for PMP22. These compounds have variable effects on the ordered partitioning of PMP22. VU0615562 and VU0619195 have little-or-no effect on the partitioning of MAL (fig. S5A), but lower P_ordered_ for PMP22 in RBL cells (fig. S1A) and possibly in HeLa cells (albeit without statistical rigor, see Fig. 4A). The third compound, primaquine diphosphate, increases P_ordered_ for both MAL (Fig. 4A and fig. S5B) and PMP22 in HeLa cells (Fig. 4A), but not in RBL cells (fig. S1A).

**Figure 4.**
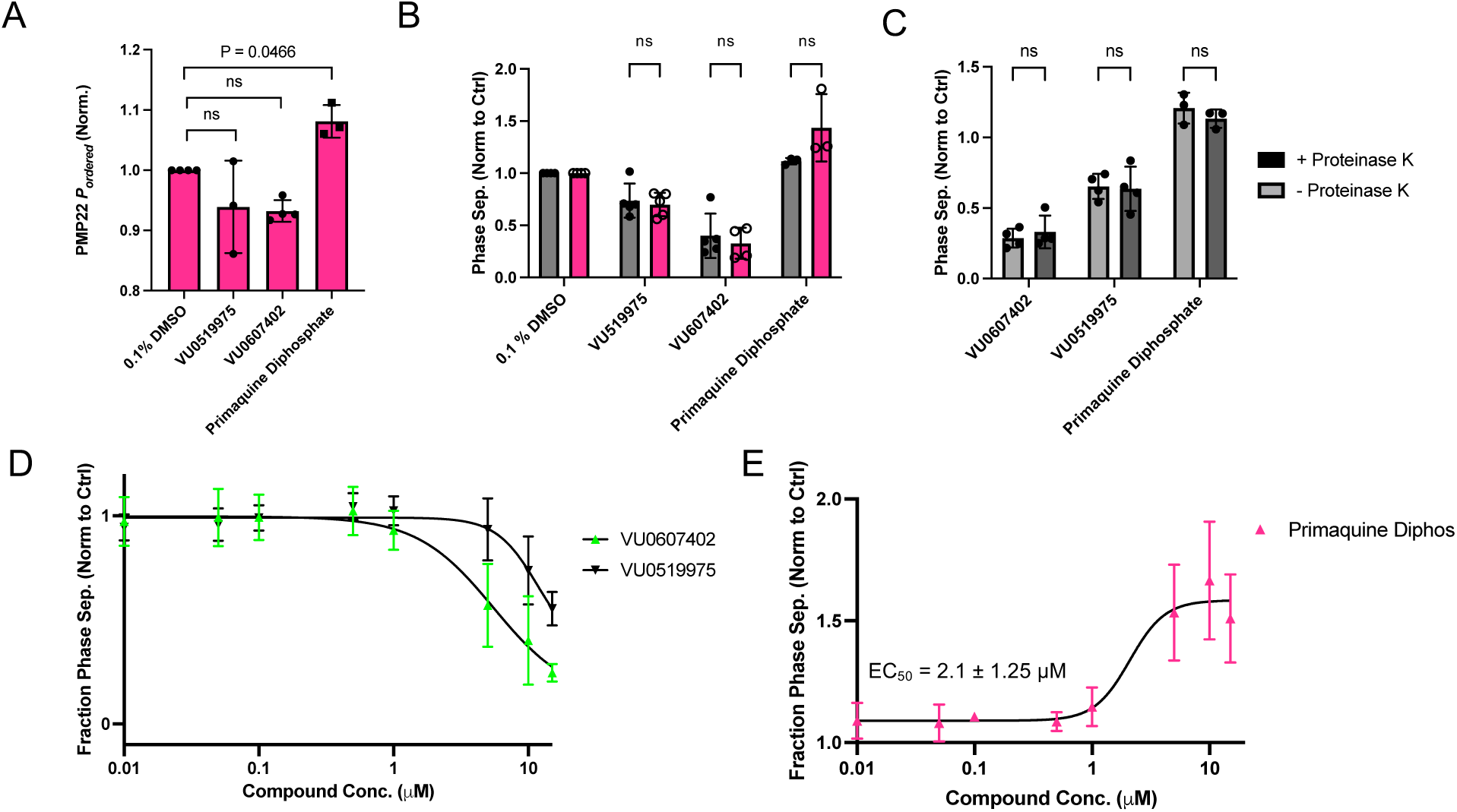
Screening identified raft modulators that are protein-independent. **A)** Effects of 10 μM VU0519975, VU06107402, and primaquine diphosphate on ordered partitioning of PMP22. Bars are means ± SD (*n* = 3), p-values are from Dunnett’s test. **B)** Effects of 10 μM VU0519975, VU06107402, and primaquine diphosphate on raft formation in GPMVs with PMP22 (magenta bars) or from untransfected cells (gray bars) PMP22 expression. Bars are means ± SD (*n* = 3-5) **C)** Effects of 10 μM VU0519975, VU06107402, and primaquine diphosphate on GPMV phase separation with (dark gray bars) and without (gray bars) proteinase K treatment **from untransfected cells (*n* = 3-5). D)** Dose response experiments with VU0519975 and VU06107402 (*n* = 3). **E)** Dose response experiments with primaquine diphosphate on raft formation. Points are means ± SD (*n* = 3). Curve and EC_50_ value from non-linear, sigmoidal fits ± SE.

Two of these compounds—VU519975 and VU607402— decreased raft formation in GPMVs (Fig. 4B, fig. S5C) while the FDA-approved drug, primaquine diphosphate (PD), increases raft formation (Fig. 4B, figs. S1B and S5D). These compounds are chemically dissimilar (Table 2). Importantly, unlike the case for Class I compounds, the presence of PMP22 (Fig. 4B) or MAL (fig. S5C) did not dramatically alter the effects of these compounds on raft formation. The independence of the activity of the compounds on proteins is supported by experiments with GPMVs from untransfected cells treated with proteinase K. The activities of these compounds were not sensitive to proteolysis (Fig. 4C). Taken together, we conclude that these compounds are raft modulators that likely interact with the membrane in a protein-independent manner. EC_50_ values for impact on raft stability in GPMVs could not be reliably determined for VU519975 and VU607402 since they were seen to be insoluble in GPMV buffer at concentrations above 15 µM (Fig. 4D), whereas the EC_50_ for the effect of PD on raft stability in GPMVs was determined to be roughly 2.1 µM (Fig. 4E).

**Table 2.**
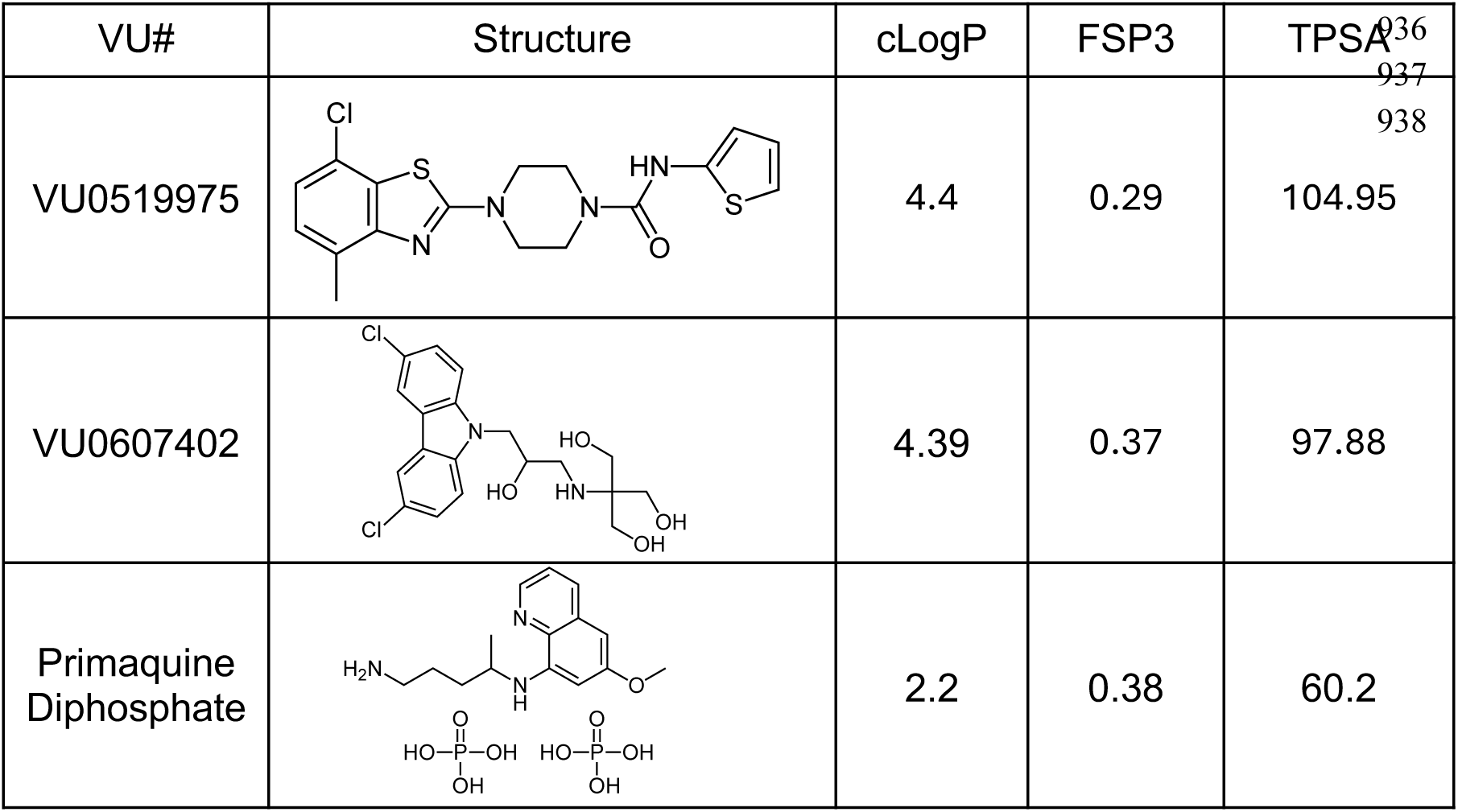
Protein-independent compound structures and parameters.

| VU# | Structure | cLogP | FSP3 | TPSA<sup>036</sup><br>937 |
| --- | --- | --- | --- | --- |
| VU0519975 |  | 4.4 | 0.29 | 104.95 |
| VU0607402 |  | 4.39 | 0.37 | 97.88 |
| Primaquine<br>Diphosphate |  | 2.2 | 0.38 | 60.2 |

Neither VU519975, VU607402, nor PD affect GPMV size or the relative size of ordered domains in GPMVs (fig. S6A-D). PD does induce a small but significant decrease in the size of PMP22-containing ordered domains (figs. S6D-bottom panel and S6E).

When interpreting any effect on raft formation it is important to factor in the impact of temperature. For example, we found that the impact of PD treatment on raft stability at 27 °C compared to 23 °C was significantly larger (a 125% increase vs a 12% increase) (fig. S5D). See Methods for additional details about this effect and considerations for data interpretation.

### Class I and II compounds alter membrane fluidity in GPMVs and live cells

We examined the effects of all five compounds (both classes, biophysical effects summarized in Table 3) on membrane fluidity to potentially shed light on how they impact on raft stability. Because fluidity experiments can be conducted in live cells this approach also provided an opportunity to connect GPMV findings with raft behavior of plasma membranes in living cells. For this, we used the environmentally sensitive dye Di-4-ANEPPDHQ (Di-4) to report on membrane fluidity (35). Increased membrane fluidity causes a red-shift of the Di-4 emission spectrum; whereas, blue-shifted spectra arise from decreased fluidity.

**Table 3.** Summary of biophysical effects of Class I and II compounds. **a.** A statistically significant decrease is clear in GPMVs RBL cells (fig. S1A), but is borderline in HeLa cells (Fig. 4A). **b.** A statistically significant increase is seen in GPMVs from HeLa cells (Fig. 4A) but not RBL cells (fig. S1A).

| Compound | Class | Impact on $P_{\text{ordered, PMP22}}$ | Impact on $P_{\text{ordered, MAL}}$ | Impact on raft formation | Impact on raft formation modified by proteins? |
| --- | --- | --- | --- | --- | --- |
| VU0615562 | I | ↓ | ↓ | ↓ | Yes |
| VU0619195 | I | ↓ | ↓ | ↓ | Yes |
| VU0519975 | II | ↓ (borderline statistical significance)<sup>a</sup> | no change | ↓ | No |
| VU0607402 | II | ↓ (borderline statistical significance)<sup>a</sup> | no change | ↓ | No |
| Primaquine Diphosphate | II | ↑<sup>b</sup> | ↑ | ↑ | No |

First, we treated GPMVs derived from HeLa cells incubated with each compound and Di-4 and then measured emission spectra. For the four raft destabilizing compounds, a pronounced red-shift from vehicle was observed in micrographs (Fig. 5A) and in measured emission spectra (Fig. 5B). To quantify these changes, generalized polarization (GP) values were determined (Fig. 5C). GP values provide a relative means of comparing spectral shifts. For Di-4 emission, lower values correspond to a more fluid (red-shifted) environment. Decreased GP values for the four raft destabilizing compounds indicate a significant increase in membrane fluidity. While there was a small increase in GP from PD treatment the change in fluidity was not statistically significant.

**Figure 5.**
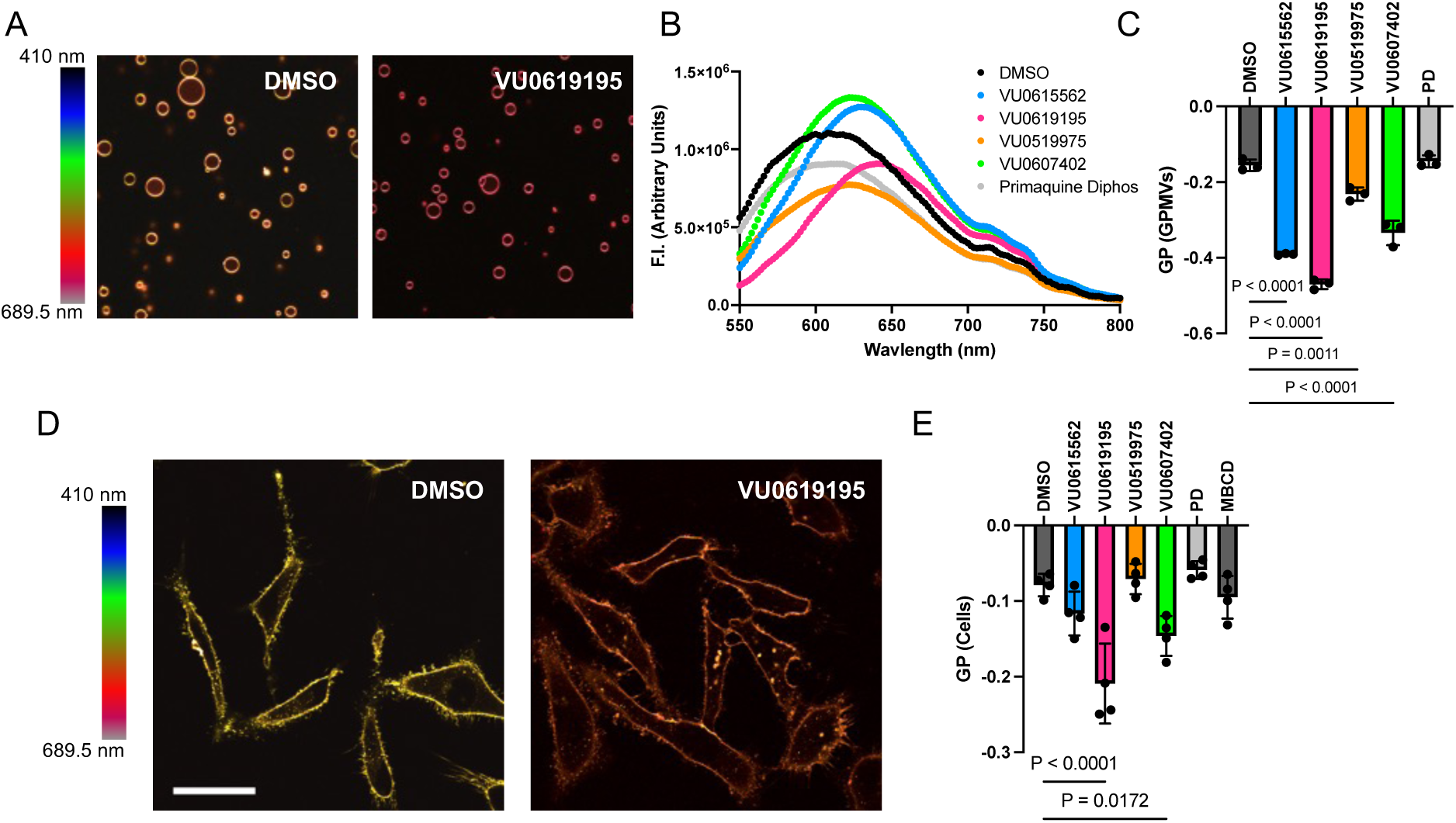
Compounds alter membrane fluidity in GPMVs and live cells. **A)** Representative spectral images (cropped from 40X fields) of GPMVs stained with Di-4 and treated with DMSO or 10 μM VU0619195. **B)** Representative Di-4 emission spectra of GPMVs treated with 10 μM hit compounds. **C)** Generalized polarization values calculated from Di-4 emission spectra shown in B. Bars are means ± SD (*n* = 3). P-values are from ANOVA followed by Dunnett’s multiple comparisons tests. **D)** Representative spectral images of live HeLa cells stained with Di-4 and treated with DMSO or 10 μM VU0619195. Scale bar = 50 μm. **E)** Generalized polarization values calculated from Di-4 emission intensities calculated from individual cells as shown in D. Bars are means ± SD, 10-15 cells (technical replicates) per treatment were measured for each of 4 biological replicates. P-values are from ANOVA followed by Dunnett’s multiple comparisons tests.

All data presented thus far were collected in GPMVs and, therefore do not inform whether the compounds exert effects in living cells. To address this, we repeated the fluidity measurements with Di-4 in live cells. HeLa cells were treated with compounds ∼15 min before a brief incubation with Di-4 to limit internalization of the dye and ensure primarily plasma membrane labeling. Cells were then imaged via laser scanning confocal microscopy using a spectral detector, allowing images from multiple emission wavelengths to be simultaneously acquired (Fig. 5D and fig. S7A. GP values were then calculated from the confocal microscopy data. We found that the changes in GP were very similar to those from GPMVs, with more variability in living cells (Fig. 5E). Two compounds, VU0619195 (Class I) and VU0607402 (Class II), produced statistically significant increases in membrane fluidity. We also added methyl-β-cyclodextrin (MBCD) to HeLa cells to deplete membranes of cholesterol and increase membrane fluidity. In comparison to addition of VU0619195 and VU0607402, the effect of MBCD treatment was modest. While not statistically significant, the effects of VU0615562, VU0519975, and PD on Di-4 emission in live cells show similar changes relative to each other as in the GPMV Di-4 experiments (Fig. 5, compare C and E).

Qualitatively, the MBCD-treated cells also appeared rounded (fig. S7, bottom row) compared to the hit compound-treated cells, which retained the well spread appearance of untreated cells, suggesting that the compounds reported here are better tolerated by live cells than MBCD treatment. Trypan blue cell viability experiments confirmed that MBCD shows toxicity in cells while none of the 5 compounds of this work were cytotoxic (fig. S7B) at the concentrations and timescales used in the live cell experiments of this study. These compounds provide improved tools for manipulating lipid rafts in living cells in studies of the functional relevance of lipid rafts in different biological processes, both by being more effective at increasing membrane fluidity and also by being less toxic. Knowing that the effects of the compounds discovered in this work extend to the plasma membranes of living cells, we were motivated to investigate their impact on putative raft-dependent functions of selected membrane proteins.

### Compounds alter TRPM8 channel activity but not autophosphorylation of EGFR

To test if Class I and Class II compounds could alter a reported lipid raft-dependent biological outcome, we examined their effects on the human TRPM8 ion channel function. TRPM8 is a cold and menthol sensing ion channel (36, 37). Many ion channels, including TRPM8, are thought to associate with lipid rafts and have signaling properties that are thought to be sensitive to being in or out of rafts (38–40). We conducted automated patch clamp (APC) electrophysiology experiments (Fig. 6A, fig. S8A) to determine if compound treatment altered TRPM8 function. TRPM8-expressing cells were equilibrated to 30 °C (41) and treated with the canonical TRPM8 agonist menthol before and after 15 minutes of continuous perfusion with the lipid raft modulating compounds. Comparisons of the pre- and post-menthol-stimulated currents were used to evaluate the effects of the raft-modulating compounds on TRPM8 function. As with the fluidity experiments, MBCD was included for comparison since previously published studies used MBCD to deplete cholesterol and disrupt lipid rafts. We found that perfusion with VU0619195 (Class I), VU0607402 (Class II), and MBCD reduced the TRPM8 current responses to menthol stimulation (Fig. 6B, fig. S8B). This result supports the notion that TRPM8 function is dependent on lipid rafts and/or changes in membrane fluidity. PD, (class II, increases raft stability) did not significantly alter TRPM8 activity. These results are similar to those seen in the live-cell fluidity experiments and provides important cross-validation.

**Figure 6.**
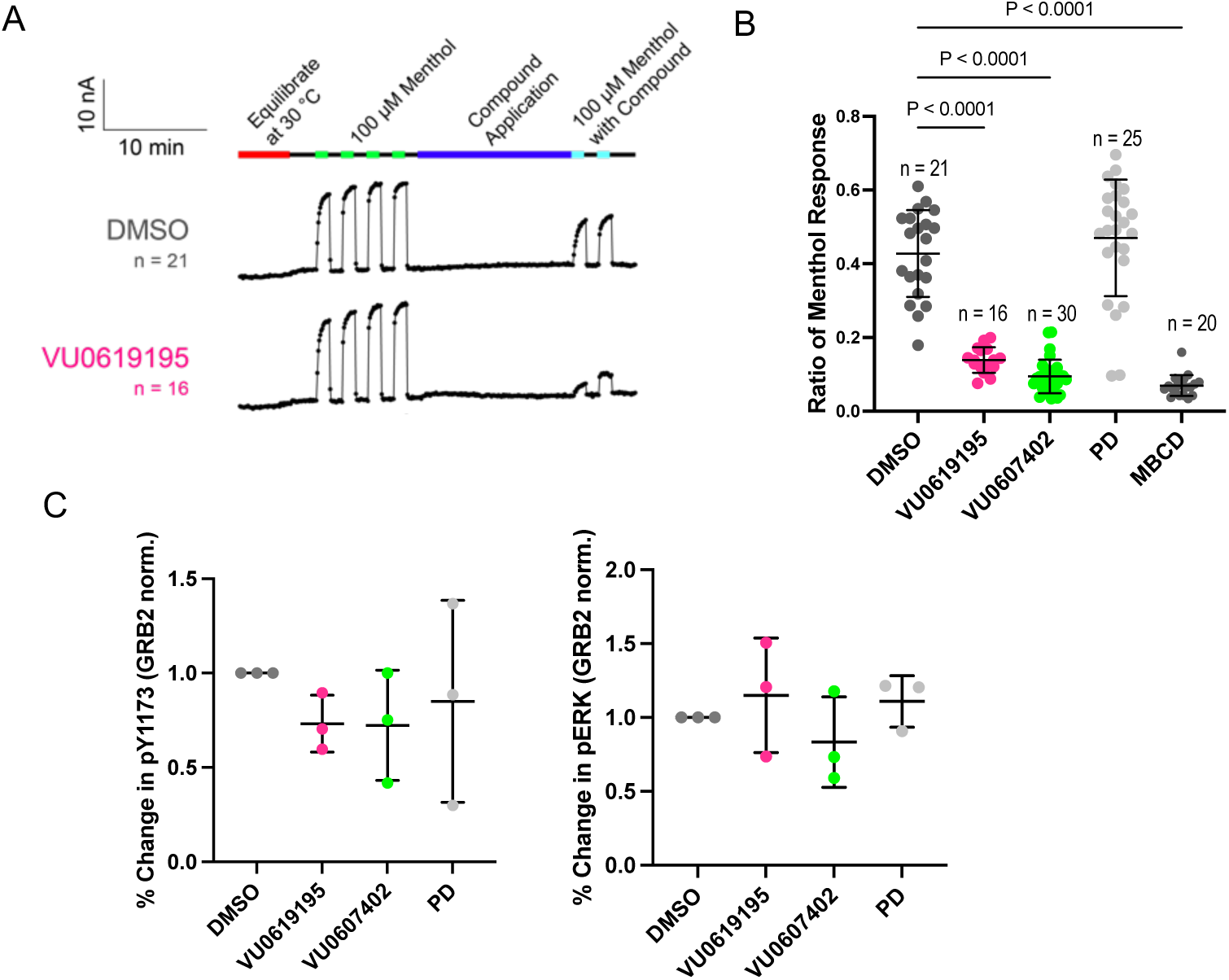
Class I and Class II compounds alter activity of TRPM8 but not EGFR. **A)** The average current traces of TRPM8 menthol response before and after exposure to the compounds in stably expressing full-length human TRPM8 HEK293T cells. Each dot is when the pulse program is applied. No perfusion was done in the first 5 minutes of the experiment to equilibrate the automated patch clamp plate to 30 °C. Menthol was applied at 100 µM for 75 seconds 4 times to allow the menthol response to saturate before perfusing continuously either 0.03% DMSO (control), 10 µM VU0619195 for 15 min (See Supp. Fig. 8A for VU0607402, PD and MBCD). 0.03% DMSO concentration was kept consistent throughout the experiment. Menthol was then applied at 100 µM with the corresponding compounds twice for 75 seconds. Each n refers to the single sum of 20 cells from an amplifier on the automated patch clamp ensemble plate. **B)** The average ratio of menthol response from each compound in stably expressing full-length human TRPM8 HEK293T cells. The ratio of menthol response uses the data from panel A and (fig. S8A), where the last two menthol response before compound application were averaged and compared to the two menthol response after compound application. Each n refers to the single sum of 20 cells from an amplifier on the automated patch clamp ensemble plate and are jittered. P-values were determined by ANOVA followed by Dunnett’s tests. Bars are means ± SD. **C)** Results of Western blot analysis of phospho-EGFR (left) and ERK (right) from HeLa cells treated with compounds for 15 min then stimulated with EGF for 1 min. n =3, bars are means ± SD. All comparisons are not significant.

The addition of VU0619195, VU0607402, and PD did not alter cell surface levels of TRPM8 as quantified by flow cytometry (fig. S8C). Interestingly, MBCD increased the level of surface TRPM8, which may be a consequence of its known activity as an inhibitor of endocytosis (42).

We also examined the impact of compounds on the activation of the epidermal growth factor receptor (EGFR) by epidermal growth factor (EGF). Previous studies have shown changes in EGFR activation following disruption of lipid rafts by MBCD treatment (43–45). We treated HeLa cells with VU0619195, VU0607402, or PD for 15 minutes (as in the TRPM8 experiments) prior to a brief (1 min) treatment with EGF. Cells were then lysed and levels of EGFR phospho-tyrosine 1173 and phospho-ERK (which is phosphorylated downstream of EGFR phosphorylation) were quantified via Western blot analysis. We did not see significant effects on either EGFR or ERK phosphorylation (Fig. 6C, fig. S9), indicating that these signaling processes are not sensitive to short-term changes in membrane fluidity or raft modulation. These data also indicate that not all plasma membrane signaling events are sensitive to the changes in lipid raft stability induced by these compounds. These EGFR data provide a useful counterexample to the findings that TRPM8 function is altered by these raft-modulating compounds.

## Discussion

HTS led to the discovery of five novel lipid raft modulating small molecule compounds falling into two classes that modulate different features of lipid raft stability. Two compounds function as protein-dependent raft destabilizers that also reduced the lipid raft affinity of both PMP22 and MAL (Class I). This a novel class of small molecules whose modality, to our knowledge, has never previously been reported. We also identified two protein-independent raft destabilizers and one protein-independent raft stabilizer that also enhance the lipid raft affinity of both PMP22 and MAL (Class II). That lipid rafts can be manipulated via protein-dependent and protein-independent interactions is notable. This suggests that there are independent lipid and protein-based stabilizing forces that work together to promote ordered domain formation. The compounds exerted effects both in isolated plasma membrane vesicles and in live-cell plasma membranes. They also altered the signaling of a raft-sensitive ion channel, TRPM8.

Commonly used methods for altering lipid rafts in cells include removal or delivery of cholesterol via MBCD and use of alcohols of varying chain lengths (46–48). These approaches each have limitations. MBCD efficiently removes cholesterol, but this is ultimately lethal to cells. MBCD is also not specific and can remove phospholipids as well as cholesterol (47). Hexadecanol and octanol can be used to decrease or increase membrane fluidity, respectively (46), but they have low miscibility in aqueous buffers and media, making them challenging to work with in cell-based assays. We also recently described bioactive compounds that promote and reduce raft formation, but the previously-reported compounds are all known to have effects on various cellular functions for which they were originally described (for example, one is a protease inhibitor) (30). Here we have presented new small molecules, two of which induce a more robust increase in membrane fluidity than MBCD, all of which were observed to be non-toxic under experimental conditions. These compounds should therefore be useful tools for manipulating lipid raft formation and raft-partitioning of membrane proteins in biophysical and cell biological studies.

Potential applications for these compounds are exemplified in our studies with TRPM8 and EGFR. Prior studies implicated lipid raft localization as having regulatory effects on the activities of both TRPM8 and EGFR (43, 44, 49). Our results showed that compounds that decrease raft formation robustly decrease TRPM8 signaling. We note that previous studies reported an *increase* in rat TRPM8 channel activity after treatment with MBCD (41, 50). The difference in phenotype is likely the result of speciation difference between the human TRPM8 used in this study compared to the rat TRPM8 used in the previous studies (51, 52). Speciation differences have also been seen in various other TRP channels, including TRPA1, TRPV3, and TRPV1 (53, 54)

Reduced raft formation induced by treatment with the same compounds that affected TRPM8 had little to no effect on EGFR phosphorylation, in contrast to the reported impact of lipid raft disruption by MCBD treatment (43, 44, 55). This suggests that EGFR activity may be more sensitive to the cholesterol concentration in the membrane than it is to lipid rafts. Examples of membrane proteins that sense and are regulated by lipids are replete in the literature. Some G protein-coupled receptors are known to be allosterically regulated by direct binding of cholesterol (56). Another example is provided by prior work on the yeast transcriptional regulator Mga2, which demonstrated that some proteins sense acyl chain composition rather than overall fluidity and respond with a rotational conformational change that regulates their activity (57).

Further studies will need to be conducted to establish the exact mechanisms by which the five compounds reported in this work exert their effects. The data suggest that the two Class I protein-dependent compounds work to change protein raft affinity and phase separation by altering protein-lipid interactions in a manner that may be partly but incompletely coupled to changes in membrane fluidity. They may intercalate into the membrane at protein lipid-interfaces and disrupt raft promoting interactions. The discovery of these compounds supports previous assertions that proteins play a critical role in regulating lipid rafts (58, 59). The three Class II protein-independent compounds likely work by inserting into the membrane and altering lipid packing and changing membrane fluidity (60, 61). We have no reason to suspect that these compounds change the lipid composition (as MBCD does) on the short time scales (15 minutes) they were tested, and given their structures. However, future lipidomic analyses will test this possibility. We speculate that a long-term incubation of live cells with the compounds could result in some level of membrane remodeling as cells respond to the changes in raft stability and membrane fluidity. Cells have adapted mechanisms to sense and respond to these changes that result from natural temperature changes which are believed to keep their membranes near the miscibility critical point (62–65). Further experiments with the compounds can broaden our understanding of these mechanisms.

Finally, we note that two of the Class II compounds altered the phase partitioning of PMP22 in GPMVs derived from at least some cell types, while P_ordered_ for MAL remained unperturbed (see Table 3 for summary). While this is a very preliminary result and will require further exploration and testing, it does suggest that it should be possible to discover modulators of raft affinity that are protein-specific. This would be a very welcome development both for studies of how the function of specific proteins are altered by raft association and potentially even for therapeutic applications.

Overall, the discovery of first in class molecules in this work present new tools for interrogating lipid rafts and raft proteins in cells. Moreover, our initial results using these compounds shed light on the fact that both protein-lipid and lipid-lipid interactions combine to stabilize lipid rafts and that these can be independently manipulated.

## Materials and Methods

### Cell culture

HeLa and RBL-2H3 cells and were acquired from the American Tissue Culture Collection (ATCC, Manassas Va, cat #CCL-2 and CRL-2256). T-REx-293 cells were acquired from Invitrogen (Cat # R71007). Cells were grown at 37 °C in 5% CO_2_ in a humidified incubator. HeLa cells were cultured in low glucose DMEM (Gibco # 11885084) supplemented with 10% fetal bovine serum (FBS, Gibco, #26140-079) and 1% penicillin/streptomycin (P/S, Gibco, #15140-122). RBL-2H# cells were cultured in MEM (Gibco 11095808) with 10% FBS and 1% penicillin/streptomycin. T-Rex-293 cells were cultured in high glucose DMEM (Gibco cat # 11965092) supplemented with 10% tetracycline free fetal bovine serum (Corning cat # 35-075-CV) with 1% penicillin/streptomycin.

### GPMV formation and imaging

HeLa cells were plated at ∼1.4 x 10^6^ cells total in a 150 mm plate. For PMP22 experiments: 24 hrs later cells were transfected with 15 ug of pSF PMP22 N41Q-myc using Fugene 6 (Promega cat # E2691) following the manufacturers protocol. Cells were grown for an additional 48 hrs post transfection. In counterscreening experiments, cells were transfected with a MAL-GFP construct (gift from the Levental lab (23)). GPMVs were generated using a standard protocol (19), cells were first rinsed twice with 10 ml of inactive GPMV buffer (150 mM NaCl, 10 mM HEPES, 2 mM CaCl2, pH7.4). To label the disordered phase, cells were stained with DiIC12(3) (1,1’-Didodecyl-3,3,3’,3’-Tetramethylindocarbocyanine Perchlorate) (DiI, Invitrogen cat # D383) for 10 minutes. Cells were then rinsed twice again with inactive GPMV buffer then subsequently incubated in 8.5-10 ml of active GPMV buffer (GPMV buffer + 2 mM DTT, 25 mM formaldehyde) at 37°C for 1.5 hrs. After incubating, GPMVs in solution were collected from the dish and allowed to settle at room temperature for 1 hr. 8-9 mL of GPMVs were then collected by pipetting from neither the top nor the bottom of the tube to leave behind both floating and settled debris. A mouse anti-myc antibody (Cell Signaling, 9B11) was then added at a ratio of 1:1500 and incubated for 1 hr. This was followed by incubation with an anti-mouse antibody conjugated to AlexaFluor647 (Cell Signaling, cat # 4410) at 1:15000. GPMVs were added to multi-well plates with compounds from DMSO stocks and allowed to incubate for 1.5 hrs. Plates were then imaged on an ImageXpress Micro Confocal High Content Screening System (Molecular Devices, San Jose CA) with a Nikon 40X 0.95 NA Plan Apo Lambda objective and an Andor Zyla 4.2MP 83% QE sCMOS camera, and an 89-North LDI 5 channel laser light source.

### High-throughput screen and GPMV image analysis

A pilot screen was conducted with an FDA approved drug library (1,184 compounds) by testing the library plated in triplicate. Primaquine diphosphate (PD) was found to have significant effects on promoting PMP22 ordered partitioning and increasing raft stability. It was carried through the larger screen as a positive control. Compounds were obtained from the Vanderbilt Discovery collection at the VU HTS core. This library contains over 100,000 compounds with the first 20,000 representing the greatest structural diversity. Compounds were dispensed via a Labcyte Echo 555 into 384-well plates such that final the final screening concentration for all compounds was 10 µM. GPMVs were made and labeled as described above and added to the plates containing compounds. The first and last columns of the plates were filled with positive and negative controls. Plates were incubated for at least 1.5 hours at room temperature prior to imaging. 16 images per well were collected using an IXMC as described above. Images were then analyzed using VesA. Strictly standardized mean differences (SSMDs) were used to calculate effect sizes for the PMP22 ordered phase partition coefficient (*P_ordered_*), for every well (32, 66). The SSMDs for positive (PD) and negative (DMSO) controls were used as a benchmark to select hits from each plate. A typical cutoff for selecting hits was a SSMD value of >90% of the positive control SSMD for PMP22 ordered partitioning. These criteria resulted in a 1-3% hit rate. SSMD values for PMP22 ordered partitioning were used to pick hits. For each hit we also noted the fraction of phase-separated vesicles (also calculated by VesA). A minimum of 100 GPMVs expressing PMP22 per well was required to be included as a hit. Hits were then screened by qualitatively assessing the images from screening. Hits that showed significant visible compound precipitation or extreme changes in GPMV size/shape were discarded. After screening 23,360 compounds 267 hits were identified following the above criteria, reflecting a 1.06% hit rate. These hits were then tested in triplicate against library compound. From hits that were validated at this phase, those with the largest effects were selected and reordered from a commercial vendor (20 compounds, vendor list below).

Experiments using reordered compounds were conducted in 96-well plate with duplicate or triplicate wells (technical replicates) in each plate (biological replicate). For experiments with PMP22 or MAL, GPMVs containing the overexpressed construct were analyzed. For follow-up and dose response experiments, a minimum of 10 phase-separated GPMVs per biological replicate were required for ordered partitioning measurements (determination of *P_ordered_*). For experiments without PMP22 or MAL, (untransfected cells) all GPMVs were analyzed.

### Compound repurchasing

Compounds from the Vanderbilt Discovery Collection were reordered from Life Chemicals (VU0615562, Cat. No. F3382-6184) (VU0619195, Cat. No. F3398-2024) (VU0519975, Cat No. F5773-0110) (VU0607402, Cat. No. F3255-0148) (Niagara on the Lake, Ontario, Canada). PD was acquired from Selleckchem (Cat. No. S4237) (Houston, TX, USA).

### Dose-response experiments

GPMVs from cell expressing or not expressing PMP22 or MAL were prepared and labeled as described above. Doses of hit compounds or DMSO were made by serial dilution and deposited into wells of a 96-well plate for a final concentration from 0.01 to 15 µM. The measured fraction of phase-separated GPMVs and *P_ordered_* were normalized to DMSO controls and averaged. Curves were fit in GraphPad Prism10 and half maximal effective concentrations (EC_50_) were determined using a non-linear sigmoidal model.

### Proteinase K treatment

Protease treatment of GPMVs was conducted as previously described (67). GPMVs were made from untransfected HeLa as described above and labeled with DiD and NBD-PE. GPMVs were then separated into 2 tubes. One tube was treated with 20 µg/mL of proteinase K (Macherey-Nagel cat #740506). Proteinase K and untreated GPMVs were incubated for 45 min at 37° C. 2 mM PMSF was added to quench the proteinase K treatment. GPMVs were then added to wells in a 96-well plate containing compounds for a final concentration of 10 µM. GPMVs were imaged and analyzed as described above.

### Temperature considerations when working with GPMVs

With the following exception, all of the work presented thus far was carried about at 21-23 °C. Phase separation is highly temperature dependent and previous work found that the temperature at which half of GPMVs from HeLa cells phase separate is roughly 25 °C (25). The instrument used for most of this work is limited at low end of temperature to about 21°C. So, we could not easily probe the effect size of raft destabilizing compounds by further decreasing the temperature (which would increase phase separation). Our ability to heat samples was less restricted. In addition to the altered effects on raft stability in response to PD treatment, there was also a more modest increase in MAL ordered partitioning at 27 °C vs 23 °C (fig. S5C). This illustrates that temperature is a crucial variable to consider when interpreting the effects of these compounds on raft stability and ordered partitioning.

### Plate reader membrane fluidity assay

For fluidity measurements in GPMVs, GPMVs were made as described above without any staining prior to vesiculation. Once collected and settled, GPMVs were stained with Di-4-ANEPPDHQ (Invitrogen, cat # D36802). To optimize the concentration of each stain a pilot study was conducted with 0.1% DMSO and single compound and dye concentrations ranging from 0.5 µM to 10 µM. From this, 1 µM as determined to give the best effect sizes for Di-4. For experiments with all compounds GPMVs were treated with dye then deposited in wells of a 96-well plate with compounds (10 µM final for each compound) and incubated at room temperature for about 40 minutes. After incubation the plate was read on a SpectraMax iD3 plate reader (Molecular Devices). Data were acquired using SoftMax Pro 7 version 7.1.0 (Molecular Devices). Di-4 was excited at 470 nm and emission spectra were collected from 550 to 800 nm using 2 nm steps. The photomultiplier tube gain was set to automatic with an integration time of 140 ms.

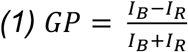

From the spectra, generalized polarization values were calculated with Equation 1. Where *I_B_* is a value in the blue end of the emission spectra and *I_R_* is a value at the red end. For Di-4 565 nm and 605 nm were selected for *I_B_* and *I_R_* respectively.

### Image-based membrane fluidity assays

For imaging, GPMVs were prepared, treated with compounds, and labeled with Di-4 in the same manner as described in the previous section. They were seeded in an 8-well chamber slide with a coverslip on the bottom (Ibidi cat # 80806). Images were acquired on a Zeiss LSM 880 laser scanning confocal using a spectral detector and a 40X oil immersion objective. Images were collected at emission wavelengths from 410 nm to 689.5 nm at an interval of 8.9 nm.

Live-cell experiments were conducted by first seeding 10,000 cells per well in 8-well chamber slides. The following day, 1 hr prior to imaging, cells treated to deplete cholesterol were first rinsed with serum-free media then incubated with 10 mM methyl-ß-cyclodextrin (MBCD) in serum-free DMEM. After 30 minutes at 37 C, cells were removed from the incubation and treated with 10 µM compounds in serum-free CO_2_ independent media (Gibco L15 media, # 2108302) for 30 min. (MBCD treated well was also swapped from 10 mM MBCD in serum-free CO_2_ independent media). Prior to imaging, Di-4 was added to a final concentration of 2 µM. Imaging and Di-4 addition were staggered to ensure less than 30 min passed after addition of the dye and imaging. This is in line with previous observations that Di-4 begins to accumulate in endosomes after 30 min. Additionally, images were acquired at room temperature to slow internalization of the dye. Imaging was conducted on the LSM 880 as described in the previous section. Fluorescence intensities of individual cells were measured in ImageJ across all 32 wavelengths. 2 to 3 cells were measured using Fiji (68) from 5 fields of view per condition. GP values were calculated as they were in the plate reader assay with intensities 561.5 nm and 605.8 nm used at the blue and red values respectively.

### Trypan blue cell viability

To ensure that effects seen in Di-4 live cell fluidity experiments and APC experiments were not due to inherent toxicity of the compounds trypan blue experiments were conducted. HeLa cells were collected and treated with 10 µM compound or 10 mM MBCD as in the Di-4 assay. Live-dead staining was then conducted with trypan blue as previously described (69). Cells were mixed in a 50:50 ratio with trypan clue reagent then immediately quantified on an automated Countess 3 cell counter (Fisher Scientific). Experiments were conducted on three separate days with measurements taken in duplicate.

### Automated patch clamp electrophysiology

HEK293 cells stably expressing full-length human TRPM8 were grown in DMEM media (Gibco 11960077) with 10% fetal bovine serum (Gibco 16000), 4 mM L-glutamine (Gibco 25030), 100 U/mL penicillin-streptomycin (Gibco 15140), 100 µM non-essential amino acid solution (Gibco 11140050), 4 mM glutaMax (Gibco 35050061), 200 µg/mL G418 (Sigma-Aldrich A1720), and 0.12% sodium bicarbonate (Gibco 25080094) at 37 °C and 8% CO_2_ in 100 mm dishes, as previously (70). After growing to 75% confluency (3-4 days), the cells were washed twice with 2 mL per dish of phosphate buffer saline solution (PBS), pH 7.4 (Gibco 10010031) followed by incubation of accutase (2 mL per dish, Gibco A1110501) for 5 minutes at 37 °C. Cells were then triturated and transferred to a conical tube and centrifuged at 200 ×g for 1.5 min to remove the accutase. The cells were resuspended with serum-free media (Gibco 11686029) and transferred into a T25 flask. The cells recovered for at least 30 mins at room temperature by gentle shaking (50 rpm). Following recovery, the cells were centrifuged (200 x g for 1.5 minutes) and resuspended in extracellular buffer (10 mM HEPES, 145 mM NaCl, 4 mM KCl, 1 mM MgCl_2_, 2 mM CaCl_2_, 10 mM glucose, pH 7.4) to a cell density of 3-7 × 10^6^ cells/mL. The osmolality of the extracellular buffer was adjusted using a Vapro 5600 vapor pressure osmometer (Wescor) with sucrose to 315-330 mOSm and pH using NaOH.

Data was collected using IonFluxMercury HT (Cell Microsystems) automated patch clamp electrophysiology instrument with Ionflux HT v5.0 software using ensemble IonFlux Plate HT (Cell Microsystems 910-0055). The ensemble microfluidic plates enable 32 parallel experiments with aggregate currents from 20 cells per experiment. Intracellular solution was composed of 10 mM HEPES, 120 mM KCl, 1.75 mM MgCl_2_, 5.374 mM CaCl_2_, 10 mM EGTA, 4 mM NaATP, pH 7.2. The intracellular solution osmolality was adjusted with sucrose to 305-315 mOsm and the pH was adjusted using KOH. Class I or II compounds were dissolved into DMSO before adding to extracellular solution, where the DMSO concentration was kept consistent at 0.03% v/v across all compounds and controls. Prior to experiments the plates were washed as suggested by the manufacturer. The protocol for the experiment is divided into four steps: prime (priming microfluidics with solutions), trap (trap cells and obtain membrane seals), break (access to intracellular by breaking membrane), and data acquisition. Each step also has multiple channels: main channel (positive pressure allows solutions to flow towards the cells and waste), trap channel (negative pressure to provide a vacuum to keep the cells in the traps and break the cell membrane to access intracellular), and compound channel (positive pressure to flow compounds to main channel). During the prime step: (1) the main channel was applied 1 psi for t = 0-25 s and 0.4 psi for t = 25-60 s (2) the trap and compound channels were applied at 5 psi for t = 0-20 s and then 1.5 psi for t = 20-55 s followed by only the traps at 2 psi for t = 55-60 s. For the voltage during the prime step, a pulse was applied every 150 ms, where the 0 mV holding potential was applied during t = 0-50 ms, 20 mV was applied during the t = 50-100 ms, and 0 mV during the t = 100-150 ms. During the trap step: (1) the main channel is applied 0.1 psi for t = 0-5 s before applying 0.5 s pulses of 0.2 psi every 5 s during t = 5-135 s (2) the trap channel is applied 6 inHg for t = 0-135 s. For the voltage during the trap step, a pulse was applied every 70 ms, where the −80 mV holding potential was applied between t = 0-20 ms, −100 mV for t = 20-50 ms, −80 mV for t = 50-70 ms. During the break step: (1) the main channel is applied 0.1 psi for t = 0-100 s (2) the trap channel was applied 6 inHg between t = 0-10 s, vacuum ramp from 10 to 14 inHg from t = 10-40 s, and 6 inHg for t = 40-100 s. For the voltage during the break step, a pulse was applied every 150 ms, where −80 mV holding potential was applied between t = 0-50 ms, −100 mV for t = 50-100 ms, and −80 mV for t = 100-150 ms. During the data acquisition: (1) the main channel is applied at 0.15 psi for t = 0-1350 s and 0 psi for 1350-2450 s, (2) the traps channel is applied 5 inHg for t = 0-3 s, 3 inHg for t = 3-1350 s, and 0 inHG for t = 1350-2450 s. For the voltage during the data acquisition, a pulse was applied every 625 ms, where the −60 mV holding potential was applied between t = 0-100 ms, −70 mV for t = 100-200 ms, −60 mV for t = 200-300 ms, a voltage ramp from −120 mV to 160 mV for t = 300-525 ms and −60 mV for t = 525-625 ms. The cells/plates were equilibrated to 30 °C for 5 minutes. Prior to application of a Class I or Class II compound, menthol was perfused for 75 s four times to measure initial current responses. Compounds were then applied by continuous perfusion for 15 minutes followed by measurement of two 75 s applications of menthol in in the presence of compound.

Ionflux Data Analyzer v5.0 was used to analyze the data. Leak subtraction was performed on the data based on the −60 mV initial holding potential and −70 mV voltage steps from the data acquisition. Each point of the current trace is from the difference of the current at 120 mV and the holding potential at −60 mV. The data was averaged from 7 points after 25 s of perfusion of menthol without or with the compound. The last two menthol stimulated currents before compound application were averaged and compared to the two menthol-stimulated currents after compound application to determine a ratio of menthol response.

### TRPM8 cell surface measurements

TRPM8 stable cells were cultured as described above. Cells were collected by dissociation with 0.5 mM EDTA in PBS and resuspended in media. Cells were then incubated in 100 µl of media with 10 µM compound or 10 mM MBCD for 15 minutes as in the electrophysiology experiments. Cells were then fixed with 100 µl Buffer A from a Fix & Perm kit for flow cytometry (Invitrogen, Cat. No. GAS004). Cells were then rinsed 3 times in flow cytometry buffer (PBS + 5% FBS + 0.1% NaN_3_). Cells were then labeled with either of two TRPM8 primary antibodies targeted to an extracellular epitope (Alomone, Cat. No. ACC-049 Abcepta, Cat. No. AP8181D). The Alomone antibody was used at a dilution of 1:100 while the Abcepta antibody was used at a dilution of 1:50 for 1 hr in the flow cytometry buffer. Cells were rinsed 3 times again then labeled with an anti-Rabbit-AlexaFlour488 secondary at a 1:1000 dilution (Cell Signaling, Cat. No. 4412) for 45 min. Cells were rinsed 3× again and resuspended in a final volume of 300 µl. Single cell fluorescence intensities were measured on a BD Fortessa 5-laser analytical cytometer. Geometric means of the resulting intensity distributions were calculated in FlowJo (version 10). Statistical comparisons were made in GraphPad Prism (version 10).

### Immunoblotting to detect EGFR activation

For EGFR activation studies, adherent HeLa cells at 70 % confluency in a 60 mm dish were starved overnight (∼18 hr) with starvation media - serum free DMEM/F12 (Gibco) media supplemented with only Pen-Step. Starved cells were exposed to small molecules of interest by replacing the overnight starvation media with 4 mL of pre-warmed starvation media containing 10 µM of Class I or II molecule of interest and 0.1 % DMSO (Cell Signaling Technologies) for noted times at 37 °C. After incubation, EGFR was activated with the addition of 1 mL of pre-warmed starvation media containing 500 ng/mL EGF (R&D Systems) for 1 minute. Due to the speed of EGFR activation kinetics in HeLa cells, treated plates were then flash frozen in liquid nitrogen after media removal. Frozen plates were then placed on ice and lysed with scraping in ice-cold RIPA lysis buffer supplemented with PhosStop phosphatase inhibitor and Complete protease inhibitor (Roche). Lysates were clarified by centrifugation and subjected to immunoblotting using NuPage Novex 4 % - 12 % Bis-Tris Protein Gels (ThermoFisher Scientific). After electrophoresis, intact gels were transferred to Immoblion-P PVDF (Millipore) membranes and cut into three horizontal strips guided by Precision Plus molecular weight ladder (Bio-Rad) and incubated overnight in blocking buffer −20 mM Tris, 150 mM NaCl, 0.1 % Tween-20 (Bio-Rad) pH 7.6 (TBST) with 3 % w/v Bovine Serum Albumin Fraction V (Fisher Bioreagents). Primary rabbit antibodies against EGFR pY1173 (53A5, 3972S), phospho-p44/42 MAPK – also known as ERK 1/2 (9101S), and GRB2 (3972S) were purchased from Cell Signaling Technology and used at a dilution of 1:1000 in TBST for 1 hr at RT with gentle agitation. Goat anti-rabbit IgG (H+L) conjugated to horse radish peroxidase (ThermoFisher Scientific, 31460) with glycerol was used at a dilution of 1:5000 for 1 hr in blocking buffer with gentle agitation. Blots were detected using SuperSignal West Pico Chemiluminescent Substrate (ThermoFisher Scientific) on a LI-COR 2800 using the chemiluminescent and 700 nm channels to detect antibody and molecular weight bands signals, respectively. Chemiluminescent signal was checked for saturation and bands of interest were integrated with Image Studio (LI-COR, version 3.1).

## Acknowledgments

The MAL-GFP construct was a gift from the Levental lab (University of Virginia). Some experiments were performed in the Vanderbilt High-Throughput Screening (HTS) Core Facility with assistance provided by Corbin Whitwell. The FDA approved library was provided by the Vanderbilt CTSA and distributed by the Vanderbilt High-Throughput Screening Core Facility as was the Vanderbilt Discovery Collection. The HTS Core receives support from the Vanderbilt Institute of Chemical Biology and the Vanderbilt Ingram Cancer Center. Membrane fluidity experiments were performed in part through the use of the Vanderbilt Cell Imaging Shared Resource.

## Author contributions

Conceptualization: KMS, AKK, WDHV, CRS

Methodology: KMS, HH, JAB, AKK, WDVH, CRS

Investigation: KMS, HH, DDL, JMH, NS, AJF, TPH

Supervision: JAB, AKK, WDVH, CRS

Writing—original draft: KMS, CRS

Writing—review & editing: KMS, HH, DDL, JAB, AKK, WDVH, CRS

## Data and materials availability

All data are available in the main text or the supplementary materials.

## References

1. Heberle, F. A., and Feigenson, G. W. (2011) Phase Separation in Lipid Membranes. Cold Spring Harb Perspect Biol. 3, a004630

2. Veatch, S. L., Rogers, N., Decker, A., and Shelby, S. A. (2023) The plasma membrane as an adaptable fluid mosaic. Biochimica et Biophysica Acta (BBA) - Biomembranes. 1865, 184114

3. Shaw, T. R., Ghosh, S., and Veatch, S. L. (2020) Critical Phenomena in Plasma Membrane Organization and Function. Annu Rev Phys Chem. 72, 51

4. Rayermann, S. P., Rayermann, G. E., Cornell, C. E., Merz, A. J., and Keller, S. L. (2017) Hallmarks of Reversible Separation of Living, Unperturbed Cell Membranes into Two Liquid Phases. Biophys J. 113, 2425–2432

5. Lingwood, D., and Simons, K. (2010) Lipid rafts as a membrane-organizing principle. Science (1979). 327, 46–50

6. Staubach, S., and Hanisch, F. G. (2011) Lipid rafts: signaling and sorting platforms of cells and their roles in cancer. Expert Rev Proteomics. 8, 263–277

7. Tsui-Pierchala, B. A., Encinas, M., Milbrandt, J., and Johnson, E. M. (2002) Lipid rafts in neuronal signaling and function. Trends Neurosci. 25, 412–417

8. Mollinedo, F., and Gajate, C. (2020) Lipid rafts as signaling hubs in cancer cell survival/death and invasion: Implications in tumor progression and therapy. J Lipid Res. 61, 611–635

9. Roy, A., and Patra, S. K. (2022) Lipid Raft Facilitated Receptor Organization and Signaling: A Functional Rheostat in Embryonic Development, Stem Cell Biology and Cancer. Stem Cell Reviews and Reports 2022 19:1. 19, 2–25

10. Brown, D. A., and London, E. (1998) Functions of lipid rafts in biological membranes. Annu Rev Cell Dev Biol. 14, 111–136

11. Varshney, P., Yadav, V., and Saini, N. (2016) Lipid rafts in immune signalling: current progress and future perspective. Immunology. 149, 13–24

12. Viljetić, B., Blažetić, S., Labak, I., Ivić, V., Zjalić, M., Heffer, M., and Balog, M. (2024) Lipid Rafts: The Maestros of Normal Brain Development. Biomolecules 2024, *Vol.* 14*, Page* 362. **14**, 362

13. Mañes, S., Del Real, G., and Martínez-A, C. (2003) Pathogens: raft hijackers. Nature Reviews Immunology 2003 3:7. **3**, 557–568

14. Parton, R. G., and Richards, A. A. (2003) Lipid Rafts and Caveolae as Portals for Endocytosis: New Insights and Common Mechanisms. Traffic. 4, 724–738

15. Head, B. P., Patel, H. H., and Insel, P. A. (2014) Interaction of membrane/lipid rafts with the cytoskeleton: Impact on signaling and function: Membrane/lipid rafts, mediators of cytoskeletal arrangement and cell signaling. Biochimica et Biophysica Acta (BBA) - Biomembranes. 1838, 532–545

16. Pralle, A., Keller, P., Florin, E. L., Simons, K., and Hörber, J. K. H. (2000) Sphingolipid–Cholesterol Rafts Diffuse as Small Entities in the Plasma Membrane of Mammalian Cells. Journal of Cell Biology. 148, 997–1008

17. Bolmatov, D., Soloviov, D., Zhernenkov, M., Zav’Yalov, D., Mamontov, E., Suvorov, A., Cai, Y. Q., and Katsaras, J. (2020) Molecular Picture of the Transient Nature of Lipid Rafts. Langmuir. 36, 4887–4896

18. Janosi, L., Li, Z., Hancock, J. F., and Gorfe, A. A. (2012) Organization, dynamics, and segregation of Ras nanoclusters in membrane domains. Proc Natl Acad Sci U S A. 109, 8097–8102

19. Sezgin, E., Kaiser, H. J., Baumgart, T., Schwille, P., Simons, K., and Levental, I. (2012) Elucidating membrane structure and protein behavior using giant plasma membrane vesicles. Nat Protoc. 7, 1042–1051

20. Fridriksson, E. K., Shipkova, P. A., Sheets, E. D., Holowka, D., Baird, B., and McLafferty, F. W. (1999) Quantitative analysis of phospholipids in functionally important membrane domains from RBL-2H3 mast cells using tandem high-resolution mass spectrometry. Biochemistry. 38, 8056–8063

21. Bauer, B., Davidson, M., and Orwar, O. (2009) Proteomic Analysis of Plasma Membrane Vesicles. Angewandte Chemie. 121, 1684–1687

22. Lorent, J. H., Diaz-Rohrer, B., Lin, X., Spring, K., Gorfe, A. A., Levental, K. R., and Levental, I. (2017) Structural determinants and functional consequences of protein affinity for membrane rafts. Nature Communications 2017 8:1. **8**, 1–10

23. Castello-Serrano, I., Lorent, J. H., Ippolito, R., Levental, K. R., and Levental, I. (2020) Myelin-Associated MAL and PLP Are Unusual among Multipass Transmembrane Proteins in Preferring Ordered Membrane Domains. J Phys Chem B. 124, 5930–5939

24. Levental, I., Lingwood, D., Grzybek, M., Coskun, Ü., and Simons, K. (2010) Palmitoylation regulates raft affinity for the majority of integral raft proteins. Proc Natl Acad Sci U S A. 107, 22050–22054

25. Marinko, J. T., Kenworthy, A. K., and Sanders, C. R. (2020) Peripheral myelin protein 22 preferentially partitions into ordered phase membrane domains. Proceedings of the National Academy of Sciences. 117, 14168–14177

26. Divincenzo, C., Elzinga, C. D., Medeiros, A. C., Karbassi, I., Jones, J. R., Evans, M. C., Braastad, C. D., Bishop, C. M., Jaremko, M., Wang, Z., Liaquat, K., Hoffman, C. A., York, M. D., Batish, S. D., Lupski, J. R., and Higgins, J. J. (2014) The allelic spectrum of charcot–marie–tooth disease in over 17,000 individuals with neuropathy. Mol Genet Genomic Med. 2, 522–529

27. Li, J., Parker, B., Martyn, C., Natarajan, C., Guo, J., Li, J., Parker, B, Martyn, C, Natarajan, C., and Guo, J. (2012) The PMP22 Gene and Its Related Diseases. Molecular Neurobiology 2012 47:2. **47**, 673–698

28. Schlebach, J. P., Narayan, M., Alford, C., Mittendorf, K. F., Carter, B. D., Li, J., and Sanders, C. R. (2015) Conformational Stability and Pathogenic Misfolding of the Integral Membrane Protein PMP22. J Am Chem Soc. 137, 8758–8768

29. Stefanski, K. M., Wilkinson, M. C., and Sanders, C. R. (2024) Roles for PMP22 in Schwann cell cholesterol homeostasis in health and disease. Biochem Soc Trans. 52, 1747–1756

30. Fricke, N., Raghunathan, K., Tiwari, A., Stefanski, K. M., Balakrishnan, M., Waterson, A. G., Capone, R., Huang, H., Sanders, C. R., Bauer, J. A., and Kenworthy, A. K. (2022) High-Content Imaging Platform to Discover Chemical Modulators of Plasma Membrane Rafts. ACS Cent Sci. 8, 370–378

31. Marinko, J. T., Wright, M. T., Schlebach, J. P., Clowes, K. R., Heintzman, D. R., Plate, L., and Sanders, C. R. (2021) Glycosylation limits forward trafficking of the tetraspan membrane protein PMP22. Journal of Biological Chemistry. 10.1016/j.jbc.2021.100719

32. Zhang, X. D. (2010) Strictly Standardized Mean Difference, Standardized Mean Difference and Classical t-test for the Comparison of Two Groups. Stat Biopharm Res. 2, 292–299

33. Skinkle, A. D., Levental, K. R., and Levental, I. (2020) Cell-Derived Plasma Membrane Vesicles Are Permeable to Hydrophilic Macromolecules. Biophys J. 118, 1292–1300

34. Bajusz, D., Rácz, A., and Héberger, K. (2015) Why is Tanimoto index an appropriate choice for fingerprint-based similarity calculations? J Cheminform. 7, 1–13

35. Jin, L., Millard, A. C., Wuskell, J. P., Dong, X., Wu, D., Clark, H. A., and Loew, L. M. (2006) Characterization and Application of a New Optical Probe for Membrane Lipid Domains. Biophys J. 90, 2563–2575

36. Peier, A. M., Moqrich, A., Hergarden, A. C., Reeve, A. J., Andersson, D. A., Story, G. M., Earley, T. J., Dragoni, I., McIntyre, P., Bevan, S., and Patapoutian, A. (2002) A TRP Channel that Senses Cold Stimuli and Menthol. Cell. 108, 705–715

37. McKemy, D. D., Neuhausser, W. M., and Julius, D. (2002) Identification of a cold receptor reveals a general role for TRP channels in thermosensation. Nature 2002 416:6876. **416**, 52–58

38. Martens, J. R., O’Connell, K., and Tamkun, M. (2004) Targeting of ion channels to membrane microdomains: Localization of K V channels to lipid rafts. Trends Pharmacol Sci. 25, 16–21

39. Bobkov, D., and Semenova, S. (2022) Impact of lipid rafts on transient receptor potential channel activities. J Cell Physiol. 237, 2034–2044

40. Kimchi, O., Veatch, S. L., and Machta, B. B. (2018) Ion channels can be allosterically regulated by membrane domains near a de-mixing critical point. Journal of General Physiology. 150, 1769–1777

41. Morenilla-Palao, C., Pertusa, M., Meseguer, V., Cabedo, H., and Viana, F. (2009) Lipid Raft Segregation Modulates TRPM8 Channel Activity. Journal of Biological Chemistry. 284, 9215–9224

42. Dutta, D., and Donaldson, J. G. (2012) Search for inhibitors of endocytosis. Cell Logist. 2, 203–208

43. Lambert, S., Vind-Kezunovic, D., Karvinen, S., and Gniadecki, R. (2006) Ligand-Independent Activation of the EGFR by Lipid Raft Disruption. Journal of Investigative Dermatology. 126, 954–962

44. Irwin, M. E., Mueller, K. L., Bohin, N., Ge, Y., and Boerner, J. L. (2011) Lipid raft localization of EGFR alters the response of cancer cells to the EGFR tyrosine kinase inhibitor gefitinib. J Cell Physiol. 226, 2316–2328

45. Chen, X., and Resh, M. D. (2002) Cholesterol Depletion from the Plasma Membrane Triggers Ligand-independent Activation of the Epidermal Growth Factor Receptor. Journal of Biological Chemistry. 277, 49631–49637

46. Machta, B. B., Gray, E., Nouri, M., McCarthy, N. L. C., Gray, E. M., Miller, A. L., Brooks, N. J., and Veatch, S. L. (2016) Conditions that Stabilize Membrane Domains Also Antagonize n-Alcohol Anesthesia. Biophys J. 111, 537–545

47. Mahammad, S., and Parmryd, I. (2015) Cholesterol depletion using methyl-β-cyclodextrin. Methods Mol Biol. 1232, 91–102

48. Suresh, P., and London, E. (2022) Using cyclodextrin-induced lipid substitution to study membrane lipid and ordered membrane domain (raft) function in cells. Biochimica et Biophysica Acta (BBA) - Biomembranes. 1864, 183774

49. Morenilla-Palao, C., Pertusa, M., Meseguer, V., Cabedo, H., and Viana, F. (2009) Lipid raft segregation modulates TRPM8 channel activity. Journal of Biological Chemistry. 284, 9215–9224

50. Veliz, L. A., Toro, C. A., Vivar, J. P., Arias, L. A., Villegas, J., Castro, M. A., and Brauchi, S. (2010) Near-Membrane Dynamics and Capture of TRPM8 Channels within Transient Confinement Domains. PLoS One. 5, e13290

51. Hilton, J. K., Salehpour, T., Sisco, N. J., Rath, P., and Van Horn, W. D. (2018) Phosphoinositide-interacting regulator of TRP (PIRT) has opposing effects on human and mouse TRPM8 ion channels. Journal of Biological Chemistry. 293, 9423–9434

52. Journigan, V. B., Alarcón-Alarcón, D., Feng, Z., Wang, Y., Liang, T., Dawley, D. C., Amin, A. R. M. R., Montano, C., Van Horn, W. D., Xie, X. Q., Ferrer-Montiel, A., and Fernández-Carvajal, A. (2021) Structural and in Vitro Functional Characterization of a Menthyl TRPM8 Antagonist Indicates Species-Dependent Regulation. ACS Med Chem Lett. 12, 758–767

53. Hilton, J. K., Rath, P., Helsell, C. V. M., Beckstein, O., and Van Horn, W. D. (2015) Understanding thermosensitive transient receptor potential channels as versatile polymodal cellular sensors. Biochemistry. 54, 2401–2413

54. Garami, A., Shimansky, Y. P., Rumbus, Z., Vizin, R. C. L., Farkas, N., Hegyi, J., Szakacs, Z., Solymar, M., Csenkey, A., Chiche, D. A., Kapil, R., Kyle, D. J., Van Horn, W. D., Hegyi, P., and Romanovsky, A. A. (2020) Hyperthermia induced by transient receptor potential vanilloid-1 (TRPV1) antagonists in human clinical trials: Insights from mathematical modeling and meta-analysis. Pharmacol Ther. 208, 107474

55. Ruzzi, F., Cappello, C., Semprini, M. S., Scalambra, L., Angelicola, S., Pittino, O. M., Landuzzi, L., Palladini, A., Nanni, P., and Lollini, P. L. (2024) Lipid rafts, caveolae, and epidermal growth factor receptor family: friends or foes? Cell Commun Signal. 22, 489

56. Jakubík, J., and El-Fakahany, E. E. (2021) Allosteric Modulation of GPCRs of Class A by Cholesterol. International Journal of Molecular Sciences 2021, Vol. 22, Page 1953. **22**, 1953

57. Ballweg, S., Sezgin, E., Doktorova, M., Covino, R., Reinhard, J., Wunnicke, D., Hänelt, I., Levental, I., Hummer, G., and Ernst, R. (2020) Regulation of lipid saturation without sensing membrane fluidity. Nature Communications 2020 11:1. **11**, 1–13

58. Levental, I., Levental, K. R., and Heberle, F. A. (2020) Lipid Rafts: Controversies Resolved, Mysteries Remain. Trends Cell Biol. 30, 341–353

59. Kervin, T. A., and Overduin, M. (2024) Membranes are functionalized by a proteolipid code. BMC Biol. 22, 1–7

60. Levental, K. R., Lorent, J. H., Lin, X., Skinkle, A. D., Surma, M. A., Stockenbojer, E. A., Gorfe, A. A., and Levental, I. (2016) Polyunsaturated Lipids Regulate Membrane Domain Stability by Tuning Membrane Order. Biophys J. 110, 1800–1810

61. Mason, R. P., Jacob, R. F., Shrivastava, S., Sherratt, S. C. R., and Chattopadhyay, A. (2016) Eicosapentaenoic acid reduces membrane fluidity, inhibits cholesterol domain formation, and normalizes bilayer width in atherosclerotic-like model membranes. Biochimica et Biophysica Acta (BBA) - Biomembranes. 1858, 3131–3140

62. Ernst, R., Ejsing, C. S., and Antonny, B. (2016) Homeoviscous Adaptation and the Regulation of Membrane Lipids. J Mol Biol. 428, 4776–4791

63. Renne, M. F., and Ernst, R. (2023) Membrane homeostasis beyond fluidity: control of membrane compressibility. Trends Biochem Sci. 48, 963–977

64. Hazel, J. R. (1995) THERMAL ADAPTATION IN BIOLOGICAL MEMBRANES: Is Homeoviscous Adaptation the Explanation? AmlU. Rev. Physiol. 57, 19–42

65. Veatch, S. L., Cicuta, P., Sengupta, P., Honerkamp-Smith, A., Holowka, D., and Baird, B. (2008) Critical fluctuations in plasma membrane vesicles. ACS Chem Biol. 3, 287–293

66. Zhang, X. D., Ferrer, M., Espeseth, A. S., Marine, S. D., Stec, E. M., Crackower, M. A., Holder, D. J., Heyse, J. F., and Strulovici, B. (2007) The Use of Strictly Standardized Mean Difference for Hit Selection in Primary RNA Interference High-Throughput Screening Experiments. 10.1177/1087057107300646. 12, 497–509

67. Dharan, R., Goren, S., Cheppali, S. K., Shendrik, P., Brand, G., Vaknin, A., Yu, L., Kozlov, M. M., and Sorkin, R. (2022) Transmembrane proteins tetraspanin 4 and CD9 sense membrane curvature. Proc Natl Acad Sci U S A. 119, e2208993119

68. Schindelin, J., Arganda-Carreras, I., Frise, E., Kaynig, V., Longair, M., Pietzsch, T., Preibisch, S., Rueden, C., Saalfeld, S., Schmid, B., Tinevez, J. Y., White, D. J., Hartenstein, V., Eliceiri, K., Tomancak, P., and Cardona, A. (2012) Fiji: an open-source platform for biological-image analysis. Nature Methods 2012 9:7. **9**, 676–682

69. Strober, W. (1997) Trypan Blue Exclusion Test of Cell Viability. Curr Protoc Immunol. 21, A.3B.1-A.3B.2

70. Luu, D. D., Ramesh, N., Can Kazan, I., Shah, K. H., Lahiri, G., Mana, M. D., Banu Ozkan, S., and Van Horn, W. D. (2024) Evidence that the cold- and menthol-sensing functions of the human TRPM8 channel evolved separately. Sci Adv. 10, 9228

