## Supplemental Materials for "Small-Molecule Modulators of Lipid Raft Stability and Protein-Raft Partitioning"

**This File Includes:**

**Supplementary Figures S1-S9 and Tables T1-T2**

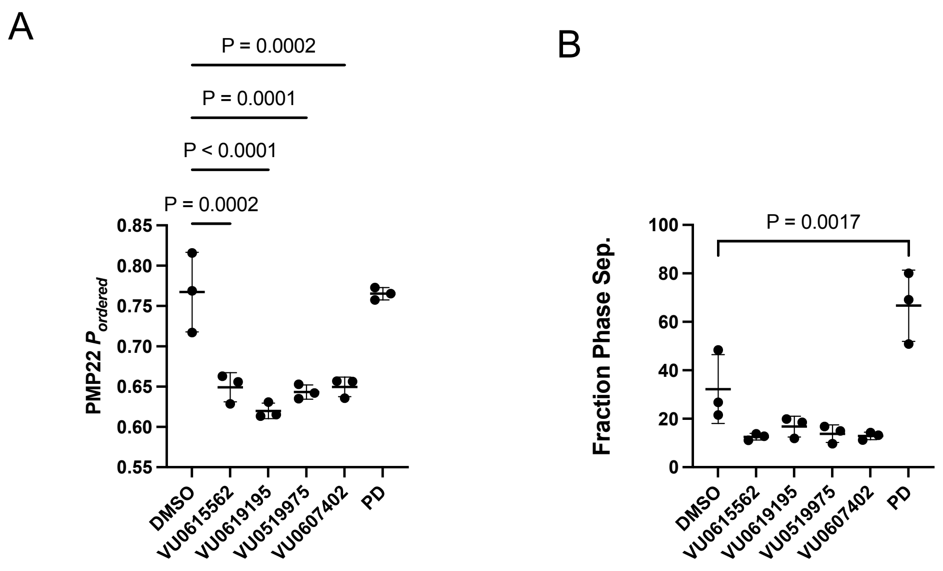


**Figure S1. Effects of Class I and II compounds on phase separation and PMP22 ordered partitioning are similar in RBL cells as compared to HeLa. A)** 5 µM ordered domain destabilizing compounds decrease decrease PMP22 ordered partitioning in GPMVs derived from RBL cells. **B)** At 5 µM, compounds that are raft destabilizing in GPMVs from HeLa cells also decrease the fraction of phase separated GPMVs from RBL cells, though the change is not statistically significant likely due to the low starting levels of phase separation in RBL GPMVs. Primaquine diphosphate significantly increases the fraction of phase separated GPMVs. n = 3, bars are means ± SD. Statistical comparisons are from ANOVA followed by Dunnett's tests.

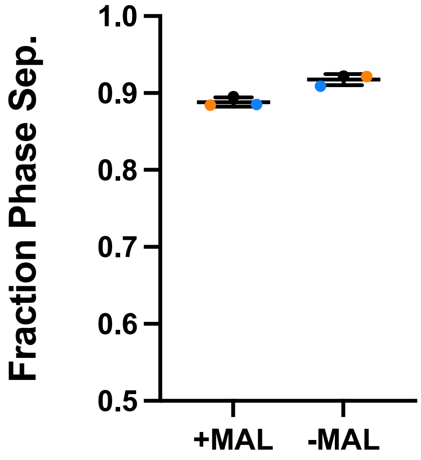


**Figure S2. MAL expression has no significant effect on raft stability.** Fraction of phase separated GPMVs from DMSO treated controls +/- MAL from the same images. Images from dose-response experiments (Fig. 2 and 3) were used. n = 3, bars are means ± SD. Individual biological replicates are shown as the same color point.

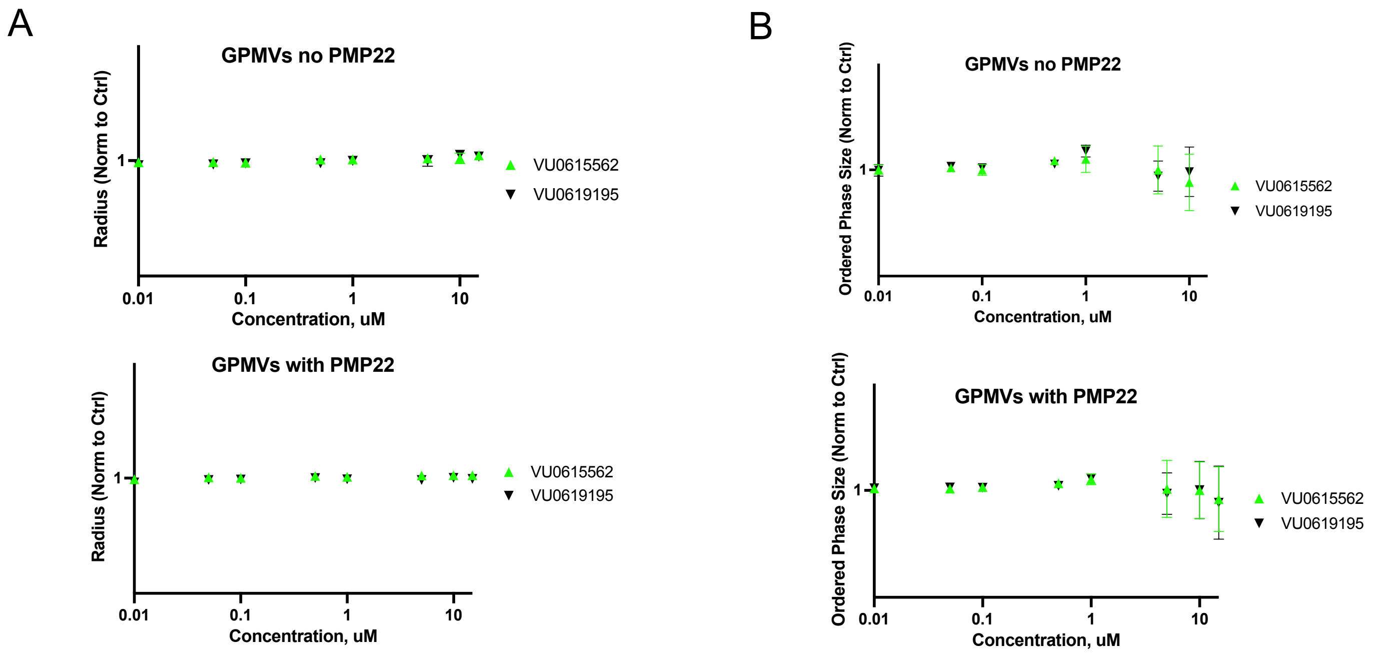


**Figure S3. Class I compounds do not affect HeLa derived GPMV size or size of ordered domains with and without PMP22 expression. A)** Average radii of GPMVs treated with VU compounds across a range of doses normalized to DMSO control. **B)** The average relative size of ordered domains in GPMVs treated with VU compounds across a range of doses normalized to DMSO control.

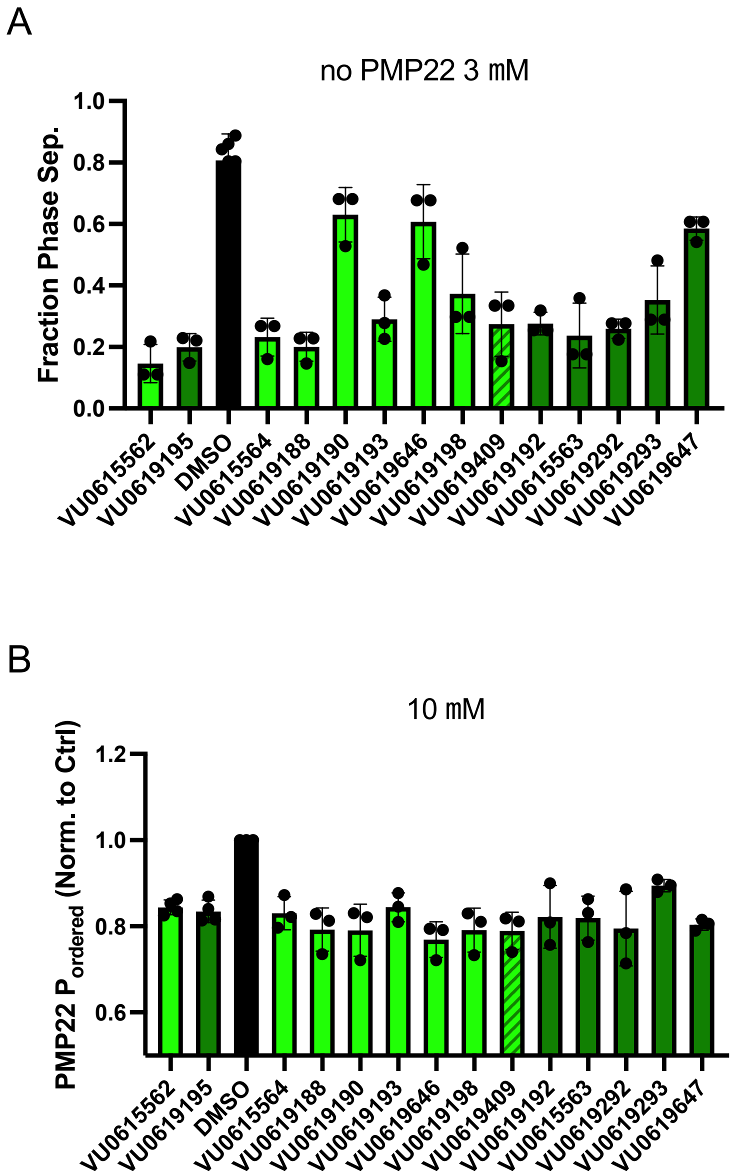


**Figure S4. SAR of protein-dependent (Class I) compounds did not identify more effective compounds, but do show that compound impact on raft formation and PMP22 partitioning are incompletely coupled. A)** Effects of SAR compounds on raft stability at 3 μM in GPMVs from untransfected cells. 3 μM was used here since effect sizes are maximal at 10 μM (the fraction of phase separated vesicles approaches 0), differences between compounds could not be statistically distinguished. Hit compounds described in the main text are shown in green bars for comparison. **B)** Effects of SAR compounds on ordered partitioning of PMP22 at 10 μM. Hit compounds VU0615562 and VU0619195 shown (green bars). $n$ = 3 bars are means ± SD. All SAR structures can be found in supplemental tables 1 and 2. Light green bars correspond to VU0615562 and associated SAR compounds (Supplementary Table 1). Dark green bars correspond to VU0619195 and associated SAR compounds (Supplementary Table 2). Hatched bar is chemically similar to both hits.

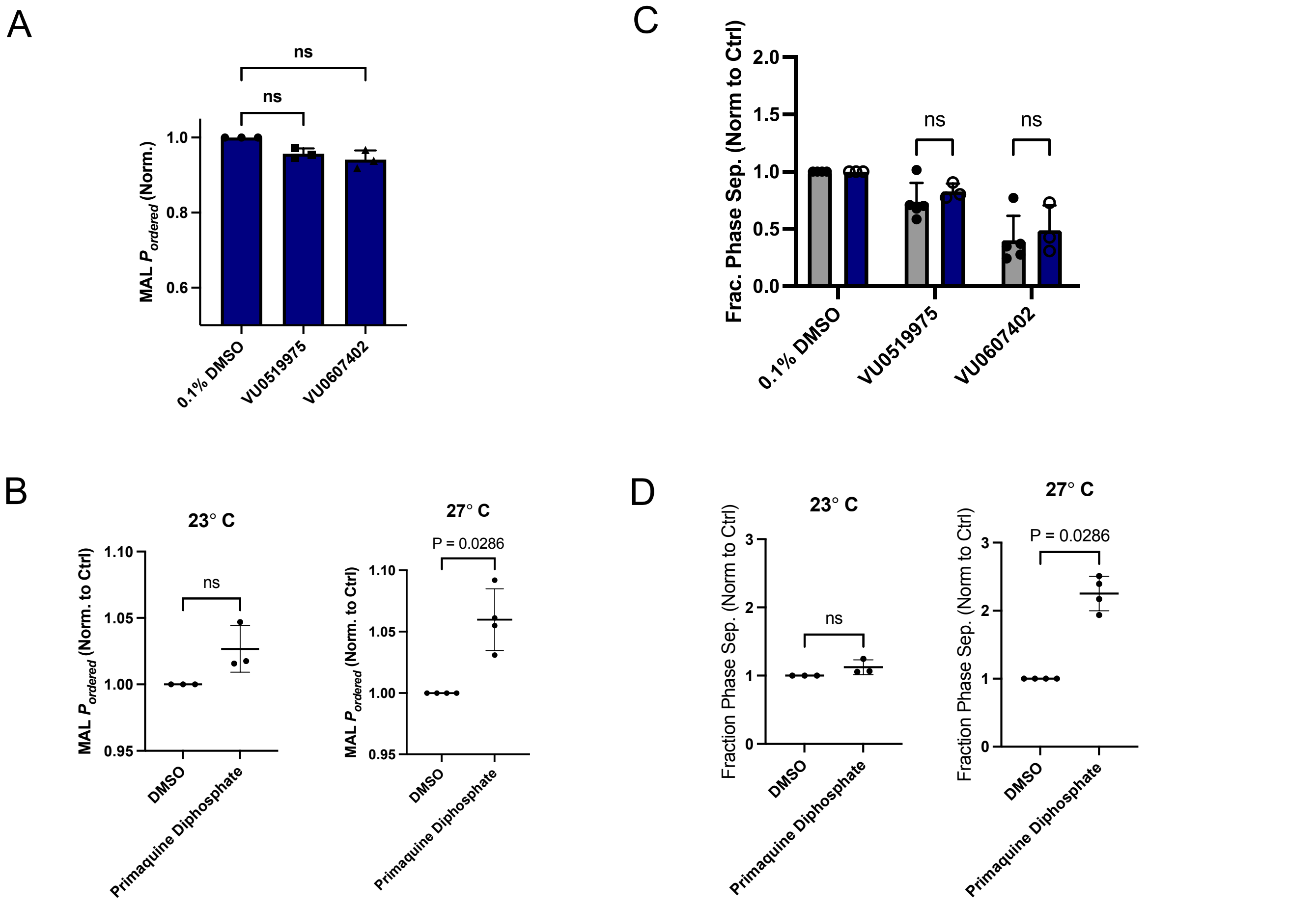


**Figure S5. Protein-independent (Class II) raft modulators have similar effects on $P_{ordered}$ and raft stability in GPMVs expressing MAL as seen with PMP22. A)** Effects of 10 µM protein-independent hits on ordered partitioning of MAL. Bars are means ± SD (*n* = 3). **B)** Effects of 10 µM protein-independent hits on phase separation with (navy bars) and without (gray bars) expression of MAL (untransfected cells). Bars are means ± SD (*n* = 3-5). **C)** Effects of 10 µM primaquine diphosphate on ordered partitioning of MAL at two temperatures. Bars are means ± SD (*n* = 3). **D)** Effects of 10 µM primaquine diphosphate on phase separation at two temperatures. Bars are means ± SD (*n* = 3-4). P-values are from Mann-Whitney tests.

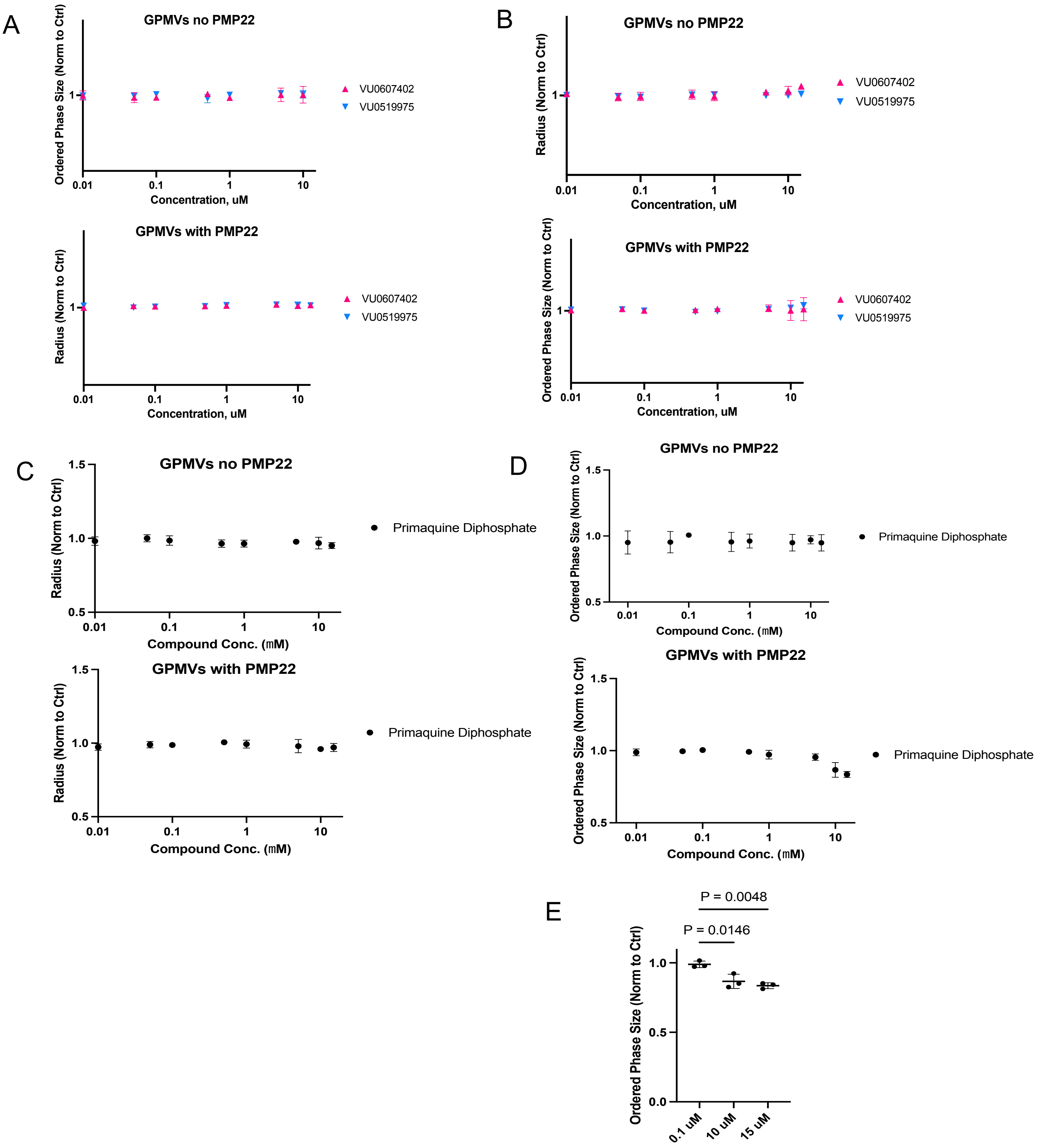


**Figure S6. Effects of Class II compounds on HeLa derived GPMV size and size of ordered domains with and without PMP22 expression. A)** Average radii of GPMVs treated with VU compounds across a range of doses normalized to DMSO control. **B)** The average relative size of ordered domains in GPMVs treated with VU compounds across a range of doses normalized to DMSO control. **C)** Average radii of GPMVs treated with primaquine diphosphate. **D)** Average ordered domain size in GPMVs treated with primaquine diphosphate. **E)** Comparison of change in domain size seen in GPMVs with PMP22 treated with PD. Points and bars are means ± SD. P-values are from ANOVA followed by Dunnetts tests.

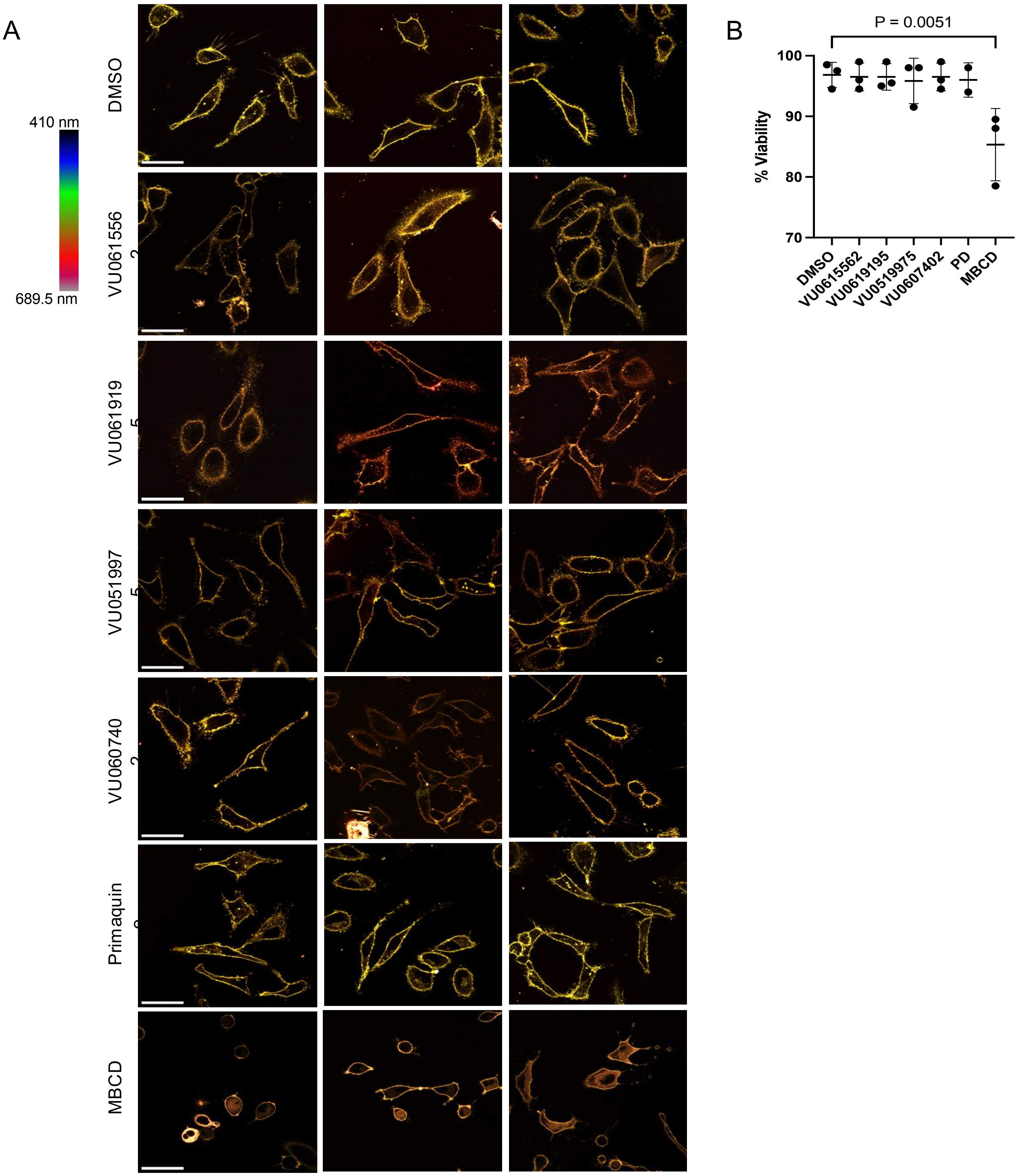


**Figure S7. Representative results of live cell membrane fluidity experiments and corresponding viability experiments. A)** Three representative spectral images from each compound treatment used in Fig 5 D and E. Cells labeled with Di-4. Scale bars are 50 μm, scale is identical for each panel. **B)** Trypan blue viability experiments conducted with HeLa cells treated with compounds as they were in the Di-4 microscopy experiments. N =3, bars are means ± SD, P-value is from ANOVA followed by Dunnett's test.

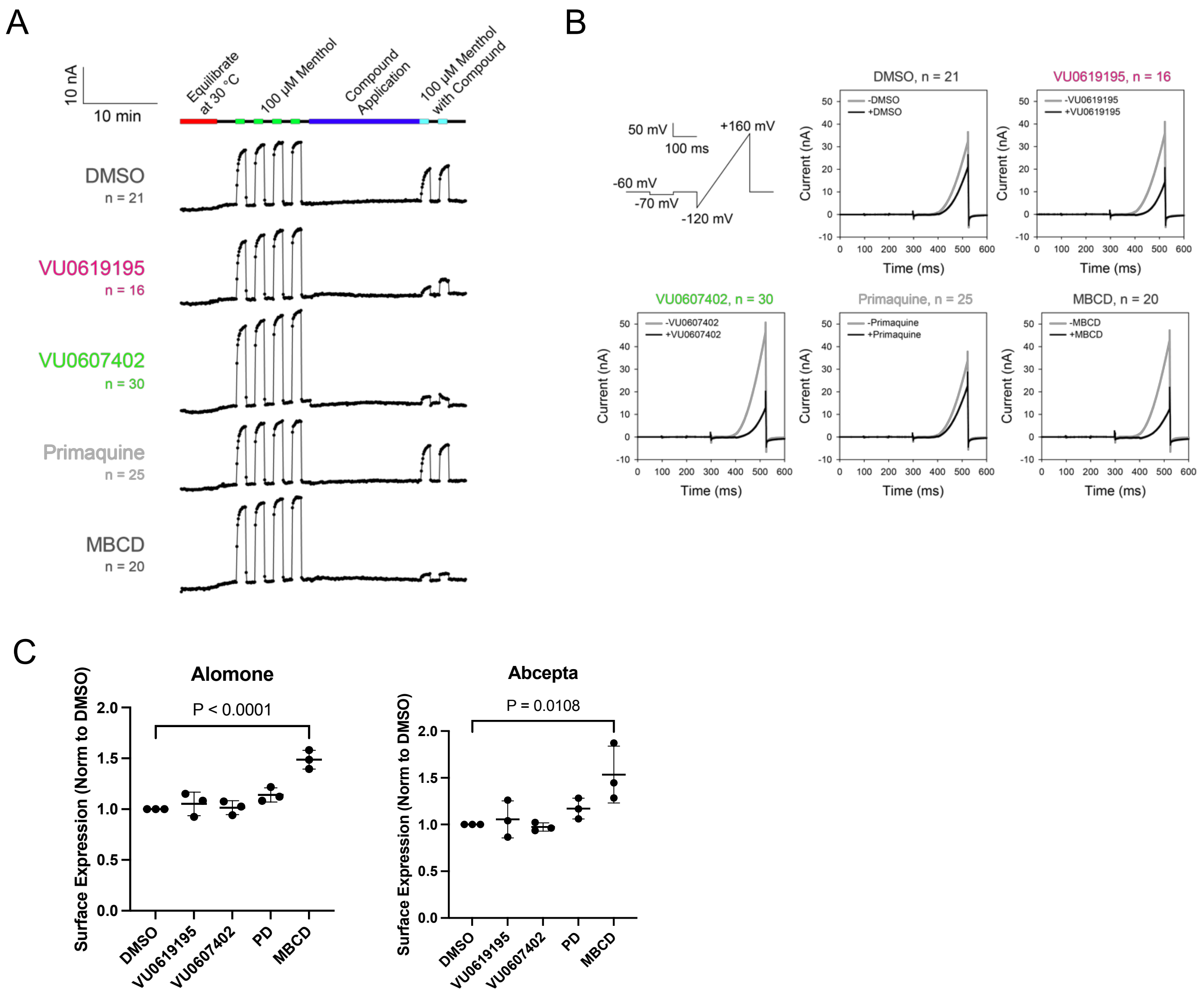


**Figure S8. Effects of Class I and Class II compounds on TRPM8 activity and cell surface expression. A)** The average current traces of TRPM8 menthol response before and after exposure to the compounds in stably expressing full-length human TRPM8 HEK293T cells. Each dot is when the pulse program is applied. No perfusion was done in the first 5 minutes of the experiment to allow the automated patch clamp plate to equilibrate to 30 °C. Menthol was applied at 100 µM for 75 seconds 4 times to allow the menthol response to saturate before applying either 0.03% DMSO (control), 10 µM VU0619195, 10 µM VU0607402, 10 µM primaquine, or 10 mM MBCD for 15 minutes. The 0.03% DMSO concentration was kept consistent throughout the experiment. Menthol was then applied at 100 µM with the corresponding compounds 2 times for 75 seconds. Each n refers to the single sum of 20 cells from an amplifier on the automated patch clamp ensemble plate **B)** The average current pulse of TRPM8 before and after exposure to the compounds. The top-left shows a schematic of the pulse program. Graphs are current response against time from a single pulse program at 922.2 seconds and 1971.99 seconds from panel A the 100 µM menthol response without (-) and with (+) 0.03% DMSO (control), 10 µM VU0619195, 10 µM VU0607402, 10 µM primaquine, or 10 mM MBCD. The 0.03% DMSO was kept consistent across all compounds. Each n refers to the single sum of 20 cells from an amplifier on the automated patch clamp ensemble plate. **C)** Quantification of plasma membrane TRPM8 in stable cells by flow cytometry using two different extracellularly directed TRPM8 antibodies following 15 min treatment with each compound as in the APC experiments. n =3 bars are means ± SD. P-values are from ANOVA followed by Dunnett's tests.

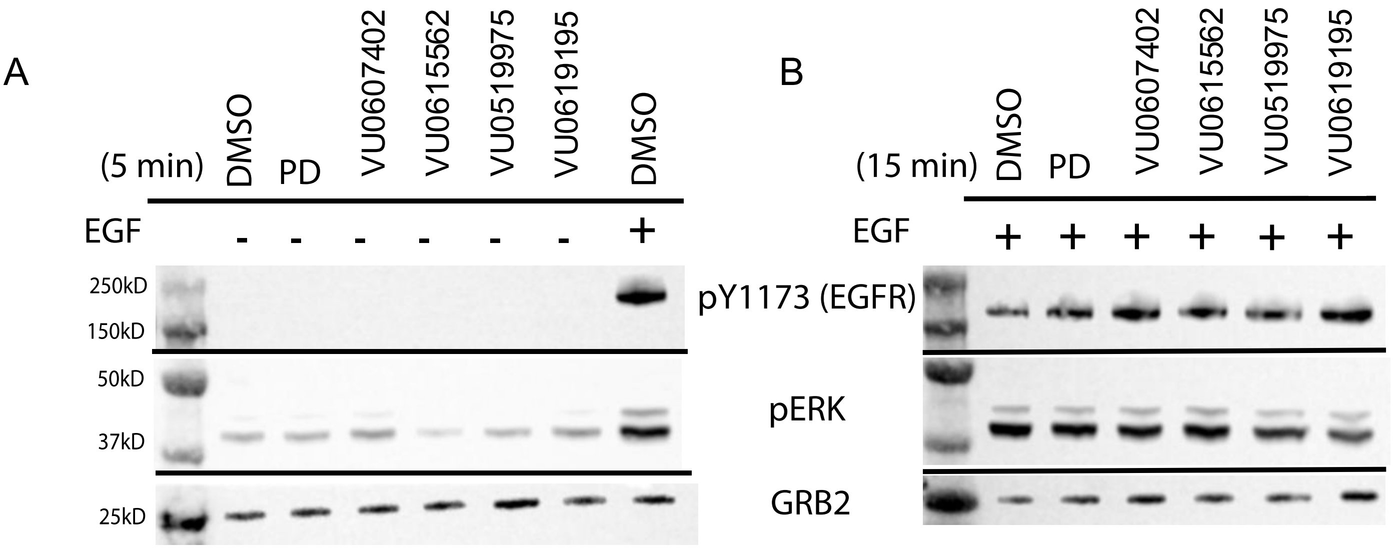


**Figure S9. Representative Western blot of phospho- EGFR and ERK. A)** Representative blot of HeLa cell lysates from cells incubated with compounds for 5 minutes and not stimulated with EGF. **B)** Representative blot of HeLa cell lysates incubated with compounds for 15 min then stimulated with EGF for 1 min.

### Table S1. VU0615562 SAR compounds

| VU ID | Structure |
|---|---|
| VU0615562 | 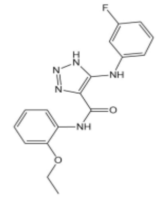 |
| VU0619188 | 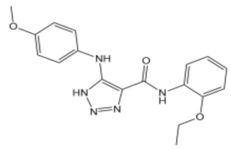 |
| VU0619190 | 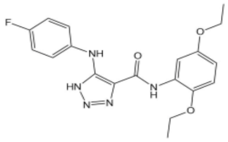 |
| VU0619198 | 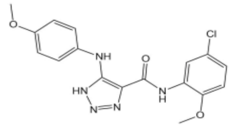 |
| VU0619409 | 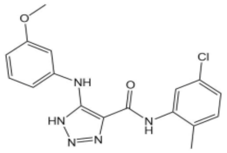 |
| VU0619193 | 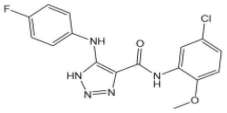 |
| VU0615564 | 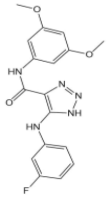 |
| VU0619646 | 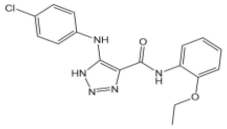 |

**Table S2. VU0619195 SAR compounds**

| VU ID | Structure |
|---|---|
| VU0615563-1 | 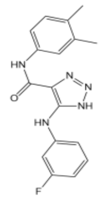 |
| VU0619192-1 | 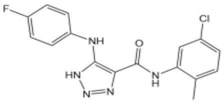 |
| VU0619195-1 | 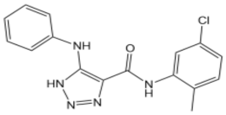 |
| VU0619292-1 | 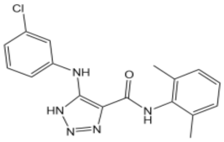 |
| VU0619293-1 | 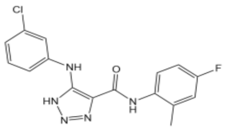 |
| VU0619409-1 | 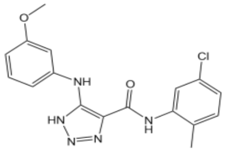 |
| VU0619647-1 | 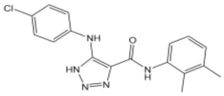 |
